# Jchain-Diphtheria Toxin Receptor Mice Allow for Diphtheria Toxin-Mediated Depletion of Antibody-Secreting Cells and Analysis of Differentiation Kinetics

**DOI:** 10.1101/2024.05.06.592703

**Authors:** KimAnh T. Pioli, Matthew Ritchie, Hira Haq, Peter D. Pioli

## Abstract

Antibody-secreting cells (ASCs) are generated following B cell activation and constitutively secrete antibodies. As such, ASCs play key roles in humoral immunity, including responses to pathogens, vaccination, and homeostatic clearance of cell debris. Therefore, understanding basic tenents of ASC biology such as their differentiation kinetics following B cell stimulation is of importance. We developed a mouse model which expresses simian heparin-binding EGF-like growth factor (*HBEGF)* [diphtheria toxin receptor (*DTR*)] under the control of the endogenous *Jchain* locus (J-DTR). ASCs from J-DTR mice expressed high levels of cell surface DTR and were acutely depleted following diphtheria toxin treatment. Furthermore, proof-of-principle experiments demonstrated the ability to use J-DTR mice to track ASC reconstitution following depletion in the spleen, bone marrow and thymus which represent sites of ASC generation and/or retention. Overall, J-DTR mice provide a new and highly effective genetic tool allowing for the study of ASC biology in a wide range of potential applications.

## Introduction

Upon stimulation, B cells have the potential to differentiate into antibody-secreting cells (ASCs)^1^ which are most prominently known for their ability to secrete thousands of antibodies (Abs) per second^2-4^. In the context of a pathogenic infection or vaccination against a specific pathogen, these Abs provide a multitude of functions (e.g., neutralization)^5^ which serve to ameliorate a current infection or act as a prophylaxis to prevent a new infection from taking hold. Therefore, it is paramount to develop a better understanding of ASC biology. This includes not only how quickly these cells are produced following B cell stimulation but also how competitive these newly formed ASCs are when confronted with limited survival niches and the presence of pre-existing ASCs. Not surprisingly, both factors have the potential to dictate the durability of the humoral immune response and seemingly are not uniform across different types of B cell subsets and stimulatory cues^6-8^.

Multiple groups have developed genetic models to fluorescently timestamp ASCs which have led to new insights regarding ASC longevity as well as the developmental relationship between short-lived plasmablasts (PBs) and post-mitotic plasma cells (PCs)^7,9-11^. Many of these experiments have been performed in the context of a fully replete ASC compartment constraining the ability to assess ASC longevity in competitive versus non-competitive scenarios. While efforts have been made to subvert these issues, these experiments largely focused on the maintenance of pre-existing ASCs. Pre-existing ASCs are critically important as they can represent a decades long record of immunization and humoral protection. However, understanding how newly formed ASCs integrate into a long-lived protective reservoir is essential especially when considering the recent COVID-19 pandemic and continual threat of newly emerging pathogens. Along these lines, being able to assess ASC generation in the presence or absence of pre-existing ASCs could be extremely informative. Towards this goal, Abs targeting CD138 have been shown to be useful in depleting ASCs in bone marrow of young and old mice^12^. However, the CD138 Ab utilized in those studies was a mouse anti-mouse Ab which included the variable region of clone 281-2^13^ and the mouse IgG2a heavy chain. This Ab was generously provided by Dr. Sherie Morrison and is not commercially available. The commercially available version of clone 281-2 was originally derived from rat splenocytes^13^ and could promote mobilization of myeloma cells (i.e., plasma cell cancer) but was unable to induce cellular depletion unless paired with bortezomib, an NF-κB inhibitor^14^. The different efficacies of these Abs may be due to the species-specific nature of the constant regions. CD138-diphtheria toxin receptor (DTR) mice have also been generated and utilized in the context of Plasmodium infection^15^. Taken together, the CD138-driven models have inherent limitations. For example, the differential level of *Cd138* expression compared to other selected immune cell types is not particularly high^11^ suggesting a narrow window allowing for ASC depletion with limited off target effects. Recently, the BICREAD mouse strain was developed which incorporates both a tamoxifen-inducible Cre recombinase (CreERT2) and *DTR* within the PR domain zinc finger protein 1 (*Prdm1)* locus^7^. While effective regarding ASC depletion, the use of *Prdm1* to drive *DTR* expression presents complications given the role of *Prdm1* in regulating both CD4 and CD8 memory T cell formation^16-18^.

To provide a tool that will allow users to modulate the presence of pre-existing ASCs, we have generated a mouse model in which the simian heparin-binding EGF-like growth factor (*HBEGF)* [*DTR*] cDNA from *Chlorocebus sabaeus* [African green monkey] was inserted into the endogenous *Jchain* locus thus targeting *DTR* expression to ASCs (referred to as J-DTR mice from here on). We have shown that these mice are functional, and that IgM, IgG and IgA ASCs can be acutely depleted following a single dose of diphtheria toxin (DT) in organs such as the spleen, bone marrow and thymus. Furthermore, due to the short half-life of DT^19^, we demonstrated the utility of this model in being able to assess ASC differentiation kinetics following depletion. At homeostasis, ASC populations were reconstituted to normal levels 7 days following DT injection supporting the concept that ASCs are continuously produced even in the absence of overt infection. Overall, the J-DTR mouse model is a highly effective tool that can be utilized to study ASC population dynamics.

## Results

### The generation of J-DTR mice and validation of DTR gene expression in ASCs

DT is derived from *Corynebacterium diphtheria* and acts as a potent protein synthesis inhibitor leading to cellular apoptosis^20^. Previous work demonstrated that DT enters the cell via receptor-mediated endocytosis following binding to the membrane bound pro-form of heparin-binding EGF-like growth factor (HB-EGF protein, encoded by the *HBEGF* gene)^21,22^. While species such as humans, simians and mice all express the HB-EGF protein, the mouse version of the protein possesses distinct amino acid differences making this species relatively insensitive to DT^23^. As such, mouse models have been developed in which human or simian *HBEGF* (referred to throughout as the *DTR*) expression is driven by cell type-specific genetic elements thus allowing for targeted ablation of that particular cell type^24-26^.

In considering an appropriate driver of *DTR* in ASCs, we examined the expression of various ASC-associated genes in the Immunological Genome Project database (ImmGen, https://www.immgen.org/) (**Figures S1A-S1C**). The ASC-associated transcription factor *Prdm1* [B lymphocyte-induced maturation protein-1 (BLIMP-1)] and cell surface marker *Sdc1* [Syndecan-1 or CD138] were readily expressed by ASC subsets; however, this was not exclusive as both genes were found to be expressed in other cell types albeit at lower levels (**Figure S1A**). Recent work utilized the *Jchain* gene to drive a CreERT2 cassette in ASCs with a great deal of success^11^. *Jchain* expression was increased in ASC populations compared to both *Prdm1* and *Sdc1* upon examination of the ImmGen data (**Figure S1B**). Furthermore, *Jchain* expression appeared highly selective for ASCs when compared to all other ImmGen cell types (**Figure S1B**) as well as those specifically in the B cell lineage (**Figure S1C**). Therefore, we generated a C57BL/6 mouse strain with the *DTR* cDNA from *Chlorocebus sabaeus* knocked into the endogenous *Jchain* locus (**Figure 1A**). In this instance, *DTR* was inserted into the *Jchain* 3’ untranslated region (UTR) downstream of an internal ribosomal entry sight (IRES) (**Figure 1A**). Upon extraction of genomic DNA, both wildtype (WT) and *DTR*-inserted *Jchain* alleles were readily identifiable by polymerase chain reaction (PCR) (**Figure 1B**).

**Figure 1:**
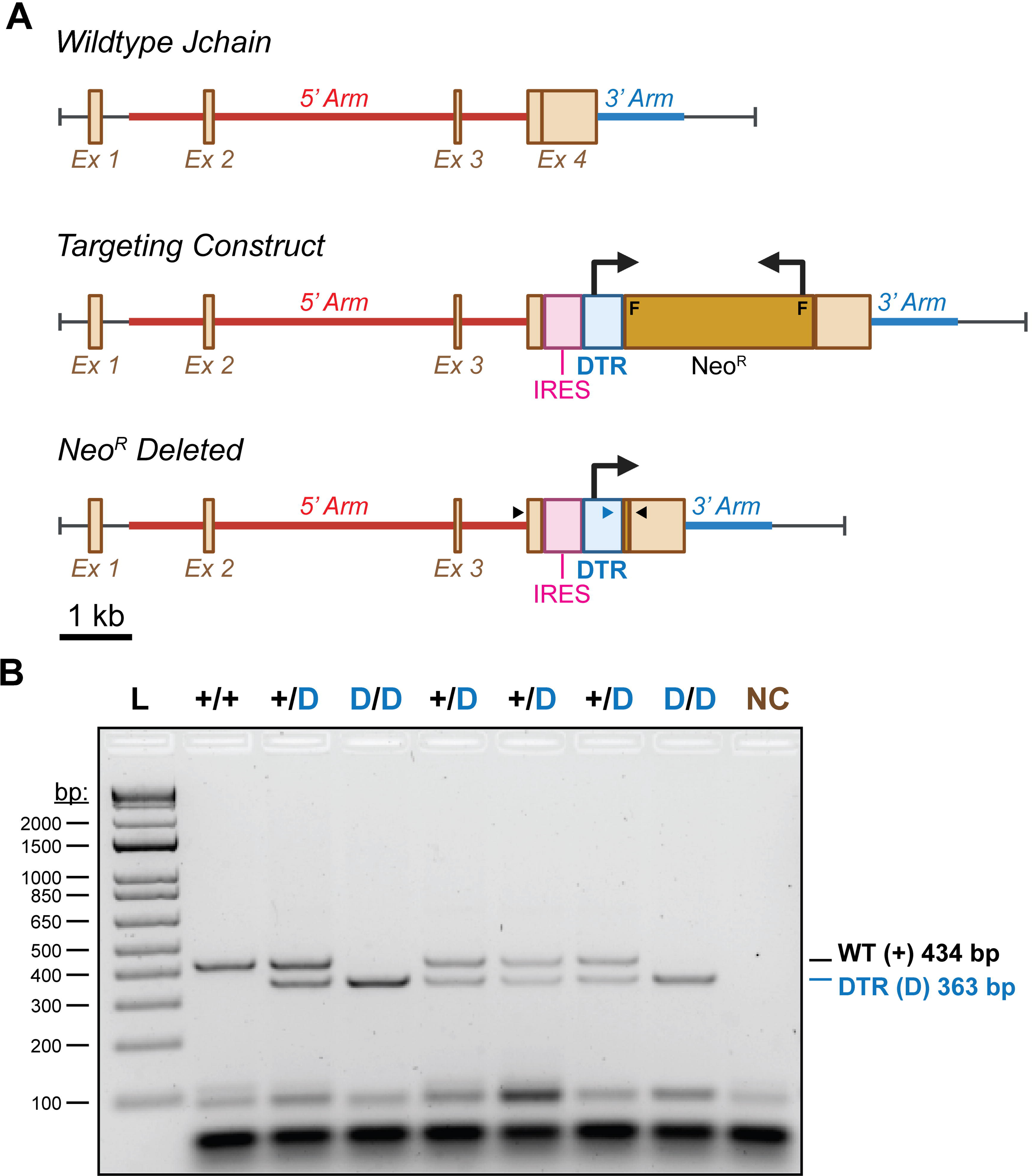
Construction and genotyping of J-DTR mice. **(A)** Schematic showing wildtype (WT) *Jchain* locus, the Targeting Construct and the final *Jchain*-*DTR* targeted insertion following Neomycin resistance (Neo^R^) cassette deletion by flippase (FLP). Schematic is drawn to approximate scale with scale bar indicating 1 kb. F = FRT sites used for FLP-mediated recombination, IRES = internal ribosomal entry site. 5’ and 3’ homology arms used to direct integration are shown. Large 90° bent arrows indicate direction of transcription for DTR and Neo^R^. Small arrow heads indicate approximate placement for genotyping primers. Black arrow heads bind DNA regions present in the endogenous *Jchain* gene. Blue arrowhead binds DNA sequence within the *DTR* coding region. Note that following Neo^R^ deletion, a single FRT site is reconstituted. This site is omitted for clarity. **(B)** Representative *Jchain-DTR* genotyping results generated from PCR amplification of genomic DNA. Note that the 434 bp WT product is only observed in animals lacking the IRES-*DTR* insertion. Animals containing this insertion would generate a product approximating 1664 bp which is not amplified using the current PCR conditions. PCR products were electrophoresed in a 2% agarose gel containing ethidium bromide. A 1 kb+ DNA ladder (L) was utilized as a size standard. NC = negative control reaction lacking DNA template.

Next, we wanted to validate that *DTR* was expressed transcriptionally in ASCs. To do so, splenocytes from female and male *Jchain*^+/+^ (WT) and *Jchain*^+/*DTR*^ (J-DTR) mice (3-7 months old) were harvested. Using Pan-B and CD138 (ASC) selection kits from STEMCELL Technologies, we enriched for splenic B cells (CD19^+^ CD138^-/LO^) and ASCs (CD138^HI^ CD267(transmembrane activator and CAML interactor,TACI)^+^) as confirmed by flow cytometry (**Figures 2A-2B and S2**). cDNA was subsequently generated from these populations and quantitative polymerase chain reaction (qPCR) analysis showed that ASC enriched samples possessed significantly higher *Prdm1* (**Figure 2C**) and *Jchain* (**Figure 2D**) gene expression. However, only ASCs enriched from the spleens of J-DTR mice showed high levels of *DTR* gene expression (**Figure 2E**). Due to variability in the effectiveness of ASC enrichment, we also examined *DTR* expression when normalized to that of *Prdm1* (**Figure 2F**). ASCs from J-DTR spleens again displayed increased levels of *DTR* transcripts when compared to WT ASCs (**Figure 2F**). In contrast, *Jchain* transcripts were equivalent between ASCs from the spleens of WT and J-DTR mice when normalized to *Prdm1* (**Figure 2G**). Western blot analysis of ASC-enriched spleen protein lysates demonstrated that insertion of *DTR* cDNA did not ablate Jchain protein expression (**Figure S3**).

**Figure 2:**
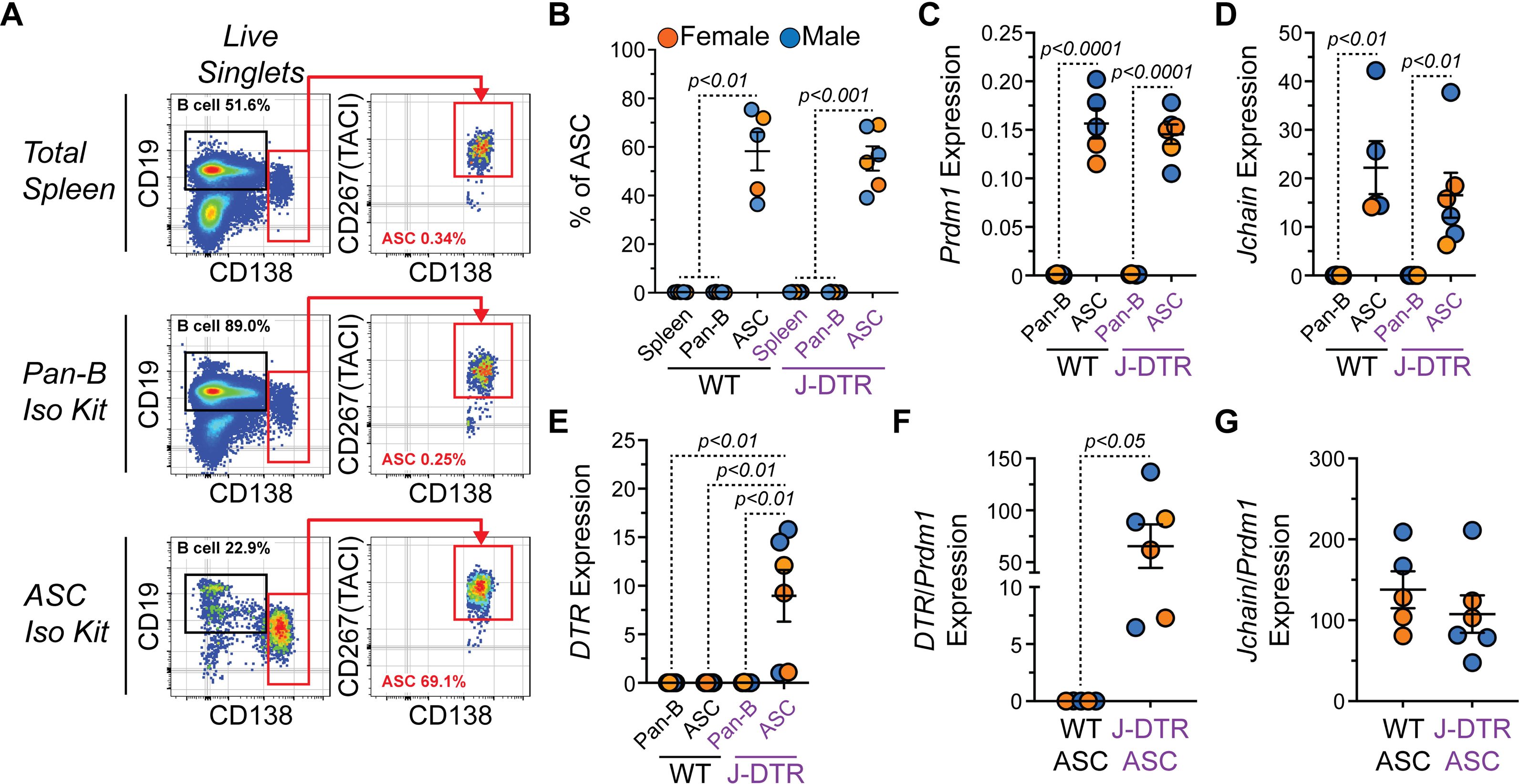
Validation of *DTR* gene expression by ASCs from J-DTR mice. **(A)** Representative flow cytometry pseudocolor plots showing gating of CD19^+^ CD138^-/LO^ B cells and CD138^HI^ CD267(TACI)^+^ ASCs in total spleen or cells purified with STEMCELL Technologies Pan-B and CD138 (ASC) isolation kits. Numbers in plots indicate percentages of gated populations within total live singlets. **(B)** Quantification of ASC percentages in total spleen or purified cells using Pan-B and ASC isolation kits. **(C-E)** Relative gene expression of **(C)** *Prdm1*, **(D)** *Jchain* and **(E)** *DTR* (*HBEGF*) in cells purified using Pan-B and ASC isolation kits. All values are relative to the expression of *Actb*. **(F)** *DTR* expression normalized to *Prdm1* expression in WT and J-DTR ASCs. **(G)** *Jchain* expression normalized to *Prdm1* expression in WT and J-DTR ASCs. **(B-G)** Symbols represent individual 3-7 months old female (orange) and male (blue) mice. Horizontal lines represent mean ± standard error of the mean (SEM). WT spleen, Pan-B and ASC: female n = 2, male n = 3; J-DTR spleen, Pan-B and ASC: female n = 3, male n = 3. **(B)** Statistics: One-way ANOVA with Dunnett’s multiple comparisons test with ASCs for each genotype set as the control column. **(C-F)** Statistics: Unpaired Student’s t-test between genotypes or cell types.

Finally, we confirmed that the DTR protein could be found on the surface of ASCs from J-DTR mice. ASCs from the spleen (**Figure 3A**), bone marrow and thymus were identified as CD138^HI^ IgD^-/LO^ CD90.2^-/LO^ CD267(TACI)^+^ CD44^+^ similar to previously published data^27^. As shown in **Figure 3B**, DTR expression was readily observable on the surface of ASCs from the J-DTR spleen compared to WT spleen ASCs or B cells (CD19^+^ CD138^-/LO^) from both genotypes. Quantification of the DTR geometric mean fluorescence intensity (gMFI), a flow cytometric readout for relative protein levels, demonstrated that J-DTR ASCs from the spleen (**Figure 3C**), bone marrow (**Figure 3D**) and thymus (**Figure 3E**) had significantly higher DTR expression compared to their WT counterparts. ASCs include 2 major populations: 1) proliferative, relatively immature and short-lived PBs and 2) post-mitotic, mature PCs with long-lived potential. In mice, CD45R(B220) expression can be used to delineate between these 2 populations (**Figure 3A**)^9,10,27,28^. In all 3 organs examined, DTR levels were increased in PBs and PCs from J-DTR mice compared to cells from WT animals (**Figures 3F-3H**). Notably, we observed increased DTR expression in PCs from the J-DTR spleen and thymus relative to the PB compartment (**Figures 3F, 3H**). Previous work has shown *Jchain* expression to be highest in IgA^+^ ASCs^29^. As such, we investigated if DTR levels were uniform amongst ASC populations expressing different Ab isotypes. Using flow cytometry, we identified membrane IgM (mIgM)^+^, membrane IgA (mIgA)^+^ and double negative (DN) ASCs in the spleen, bone marrow and thymus (**Figure S4A**). Compared to WT ASCs, all 3 ASC Ab subsets from J-DTR mice possessed increased amounts of DTR (**Figure S4B**). J-DTR mIgA^+^ ASCs had significantly higher DTR expression compared to mIgM^+^ and DN ASCs regardless of organ analyzed (**Figures S4C-S4E**). In both the spleen and thymus, mIgM^+^ ASCs from J-DTR animals displayed intermediate levels of DTR compared to mIgA^+^ and DN ASCs (**Figures S4C, S4E**).

**Figure 3:**
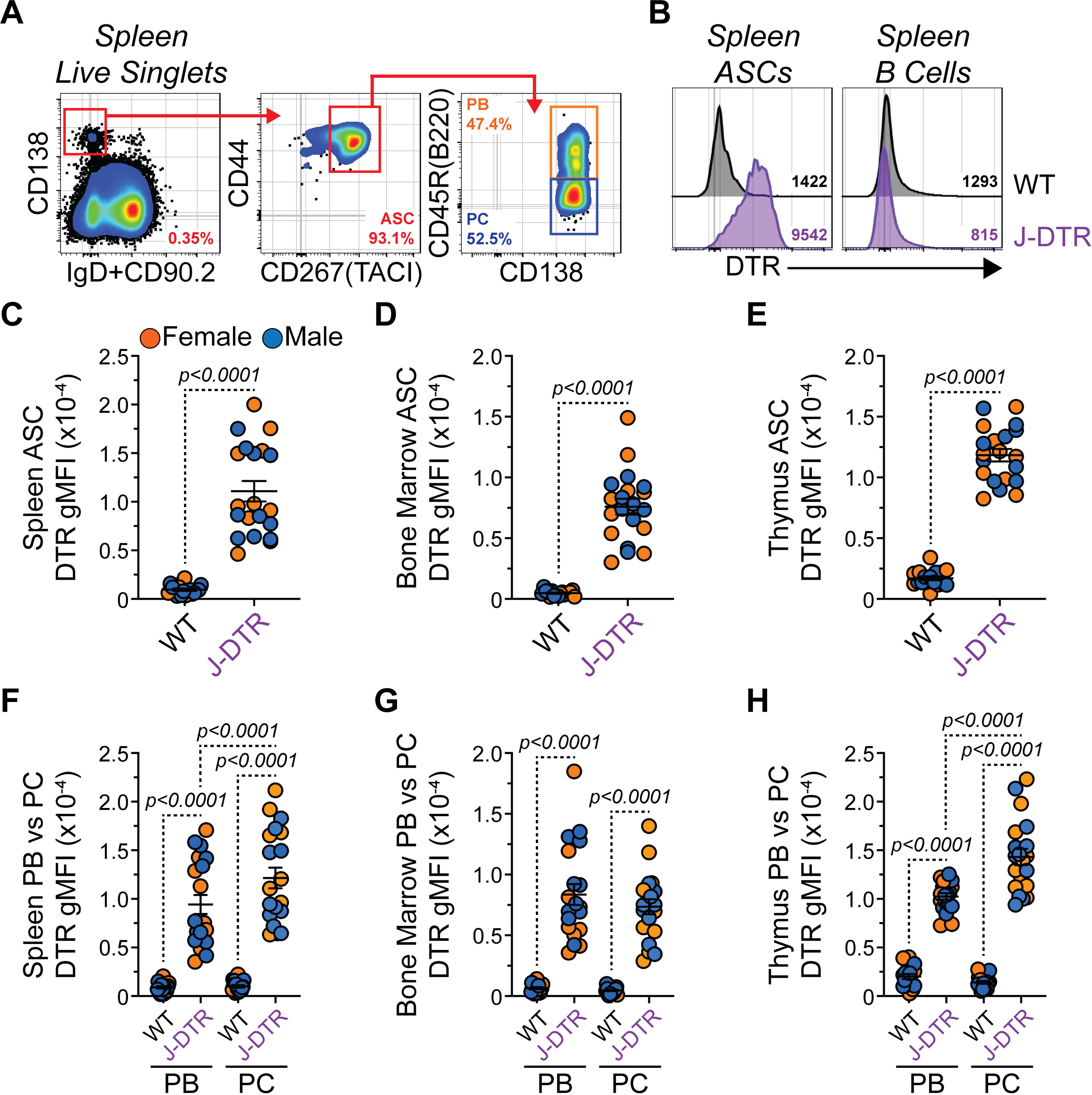
Validation of DTR surface protein expression by ASCs from J-DTR mice. **(A)** Representative flow cytometry pseudocolor plots showing gating of spleen ASCs as CD138^HI^ IgD^-/LO^ CD90.2^-/LO^ CD267(TACI)^+^ CD44^+^. ASC subset gating is shown for PBs (CD45R(B220)^+^) and PCs (CD45R(B220)^-^). Numbers in plots indicate percentages of gated populations within the immediate parent population. **(B)** Representative flow cytometry histogram overlays showing surface expression of DTR by spleen ASCs and B cells from both WT and J-DTR mice. Spleen B cells were gated as CD19^+^ CD138^-/LO^. Numbers in plots indicate DTR gMFIs. **(C-E)** gMFIs for WT and J-DTR ASCs from **(C)** spleen, **(D)** bone marrow and **(E)** thymus. **(F-H)** DTR gMFIs for WT and J-DTR PBs and PCs from **(F)** spleen, **(G)** bone marrow and **(H)** thymus. **(C-H)** Symbols represent individual 3-7 months old female (orange) and male (blue) mice. Horizontal lines represent mean ± SEM. WT: female n = 8, male n = 9; J-DTR: female n = 10, male n = 10. **(C-E)** Statistics: Unpaired Student’s t-test between genotypes. **(F-H)** Statistics: Unpaired Student’s t-test comparing between genotypes. Paired Student’s t-test comparing PBs and PCs within a genotype.

To determine if DTR was preferentially expressed in ASCs compared to other B cells types, we examined DTR surface expression on total B cells (CD19^+^ CD90.2^-^ CD138^-/LO^) as well as germinal center (or germinal center-like) B cells (GCB, CD19^+^ CD90.2^-^ CD138^-/LO^ CD95(Fas)^+^ GL7^+^) from both the spleen (**Figures S5A-S5B**) and thymus (**Figures S5C-S5D**). As a comparison within the same flow cytometry samples, we also analyzed the CD138^HI^ CD90.2^-^ compartment which would be enriched for ASCs. As expected, total B cells demonstrated minimal DTR expression while ASC-containing CD138^HI^ CD90.2^-^ cells possessed high levels of the protein (**Figures S5E-S5F**) in J-DTR animals. Within the spleen, J-DTR GCBs possessed an intermediate phenotype while the GCB-like population in the thymus seemingly lacked DTR expression (**Figures S5E-S5F**). Overall, these data suggest that while all ASCs from J-DTR mice preferentially express the DTR, some organ-specific differences may exist based upon maturation status.

### J-DTR mice demonstrate normal generation of ASCs in the spleen, bone marrow and thymus

The above studies validated DTR expression by ASCs from the spleen, bone marrow and thymus. However, it was unclear if DTR expression via insertion into the *Jchain* locus altered homeostatic generation of ASCs or secretion of various Ab isotypes. Flow cytometric quantification of total ASCs in the spleen, bone marrow and thymus showed no significant differences between WT and J-DTR mice (**Figures 4A-4C**). In terms of Ab secretion, enzyme-linked immunosorbent spot (ELISpot) assays demonstrated the presence of IgM, IgG and IgA ASCs in spleen, bone marrow and thymus (**Figure 4D**). Quantification of spot numbers for IgM (**Figure 4E**), IgG (**Figure 4F**) and IgA (**Figure 4G**) showed no significant alterations between WT and J-DTR mice in the spleen, bone marrow or thymus.

**Figure 4:**
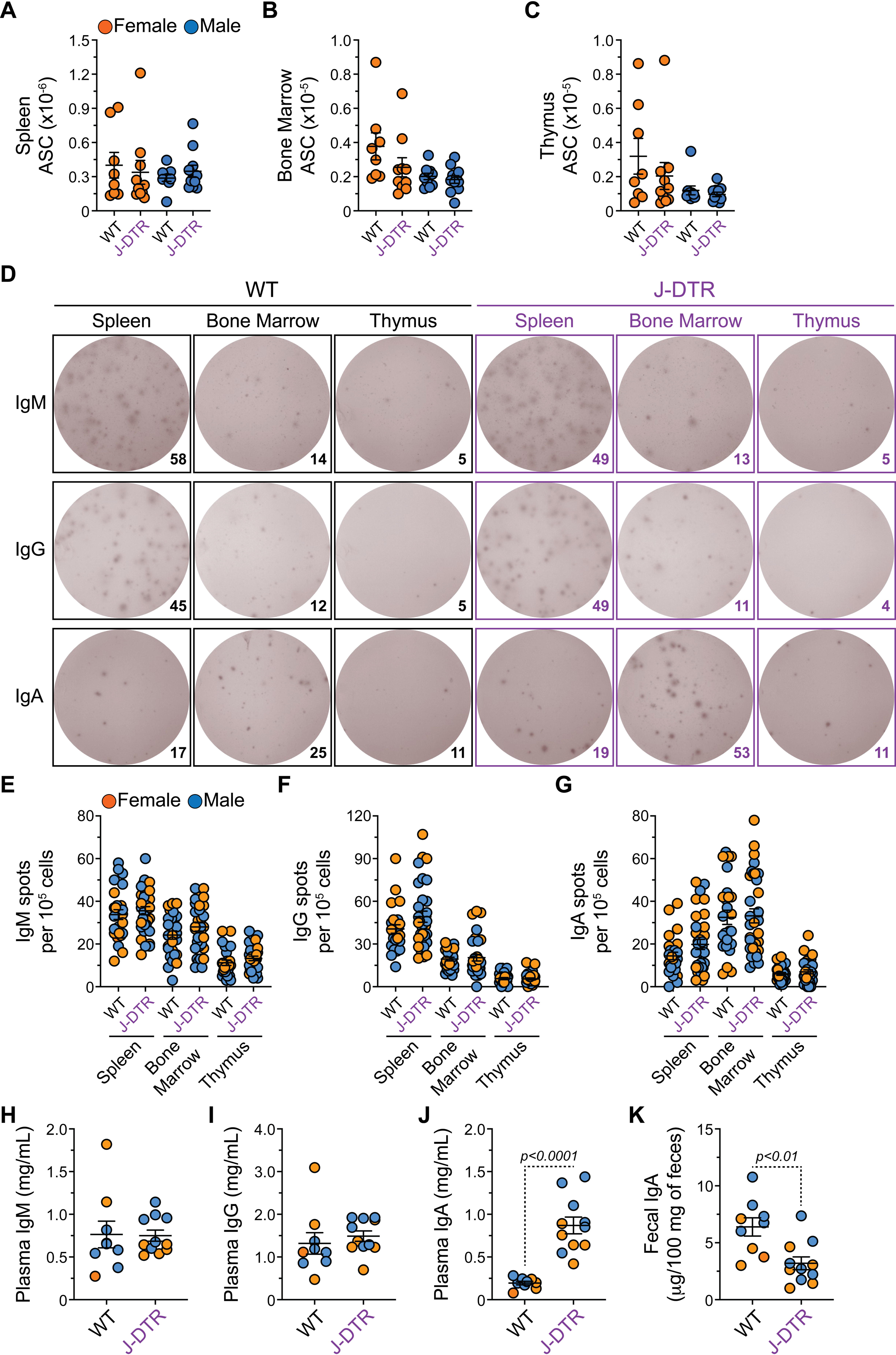
ASCs develop normally in the spleen, bone marrow and thymus of J-DTR mice. **(A-C)** Total ASC numbers for **(A)** spleen, **(B)** bone marrow and **(C)** thymus of WT and J-DTR mice. **(D)** Representative ELISpot images showing IgM, IgG and IgA spot formation from spleen, bone marrow and thymus of WT and J-DTR mice. 10^5^ cells per well were plated in triplicate. Numbers indicate spots counted per well following background subtraction of identical wells not coated with capture Ab. **(E-G)** Numbers of **(E)** IgM, **(F)** IgG and **(G)** IgA spots per 10^5^ cells from spleen, bone marrow and thymus of WT and J-DTR mice. **(H-J)** Plasma concentration of **(H)** IgM, **(I)** IgG and **(J)** IgA from WT and J-DTR mice. **(K)** Fecal IgA concentration from WT and J-DTR mice. **(A-C)** Symbols represent individual 3-7 months old female (orange) and male (blue) mice. Horizontal lines represent mean ± SEM. WT: female n = 8, male n = 10; J-DTR: female n = 10, male n = 11. Statistics: Unpaired Student’s t-test between genotypes. **(E-G)** Symbols represent individual wells plated in triplicate from 3-5 months old female (orange) and male (blue) WT and J-DTR mice. Values shown represent data following subtraction of spots detected in wells that were not coated with capture Ab (background). Horizontal lines represent mean ± SEM. WT: female n = 4, male n = 5. J-DTR: female n = 5, male n = 6. Statistics: Unpaired Student’s t-test between genotypes. **(H-K)** Symbols represent individual 3-5 months old female (orange) and male (blue) WT and J-DTR mice. Horizontal lines represent mean ± SEM. WT: female n = 4, male n = 5. J-DTR: female n = 5, male n = 6. Statistics: Unpaired Student’s t-test between genotypes.

Enzyme-linked immunosorbent assay (ELISA) assessment of plasma Ab isotypes demonstrated concordant results with plasma IgM (**Figure 4H**) and IgG (**Figure 4I**) being similar between WT and J-DTR mice. In contrast, IgA was significantly elevated in J-DTR plasma (**Figure 4J**). While IgA is relatively rare in the plasma, it is present in high amounts in the feces^30^ due to transport across the intestinal epithelium which largely requires the Jchain^31,32^. Analysis of feces revealed lower IgA amounts in the fecal supernatants isolated from J-DTR mice (**Figure 4K**). In total, ASC generation appeared normal in J-DTR animals excluding tissue-specific alterations in IgA levels.

### DT treatment leads to acute ASC depletion in J-DTR mice

To test the functionality of our J-DTR mouse model, we administered intraperitoneal (i.p.). injections of either phosphate buffered saline (PBS) or 200 ng DT (100 μL volume) to both female and male, 3-4 months old WT and J-DTR animals (**Figures 5, 6 and S6-S7**). The next day, mice were euthanized and ASC populations in the spleen, bone marrow and thymus were examined by flow cytometry (**Figures 5A-5B**). DT had no overall impact on the cellularity of any organ from either genotype (**Figures S6B-S6D**). In all 3 organs assessed, DT treatment led to significant reductions in ASC numbers only in J-DTR mice (**Figures 5C-5E**). Notably, there was some variability in this effect as spleen (**Figure 5C**), bone marrow (**Figure 5D**) and thymus (**Figure 5E**) ASCs were reduced by ∼40x, 33x and 12x, respectively. Our analysis of DTR expression indicated some differences based upon maturation status (**Figures 3F, 3H**) with PCs having higher expression than PBs in the spleen and thymus. As such, we also assessed if DT-mediated ablation differentially impacted PBs versus PCs (**Figures 5F-5H**). Independent of organ, DT injection led to a significant depletion of PBs and PCs in J-DTR mice with the overall magnitude (i.e., fold difference) being highest for PCs (**Figures 5F-5H**). This suggested that DT-mediated elimination of PCs was more efficient compared to PBs. To support this, we used PB and PC numbers to generate a PB:PC ratio for the spleen (**Figure S6E**), bone marrow (**Figure S6F**) and thymus (**Figure S6G**) of J-DTR mice that received either PBS or DT. Following DT treatment of J-DTR mice, the PB:PC ratio increased suggesting a population shift to a more immature PB phenotype (**Figures S6E-S6G**). Furthermore, we observed that residual ASCs present in DT-treated J-DTR mice expressed significantly less DTR compared to PBS-treated J-DTR animals (**Figures S6H-S6J**); however, the significance of this observation remains to be determined.

**Figure 5:**
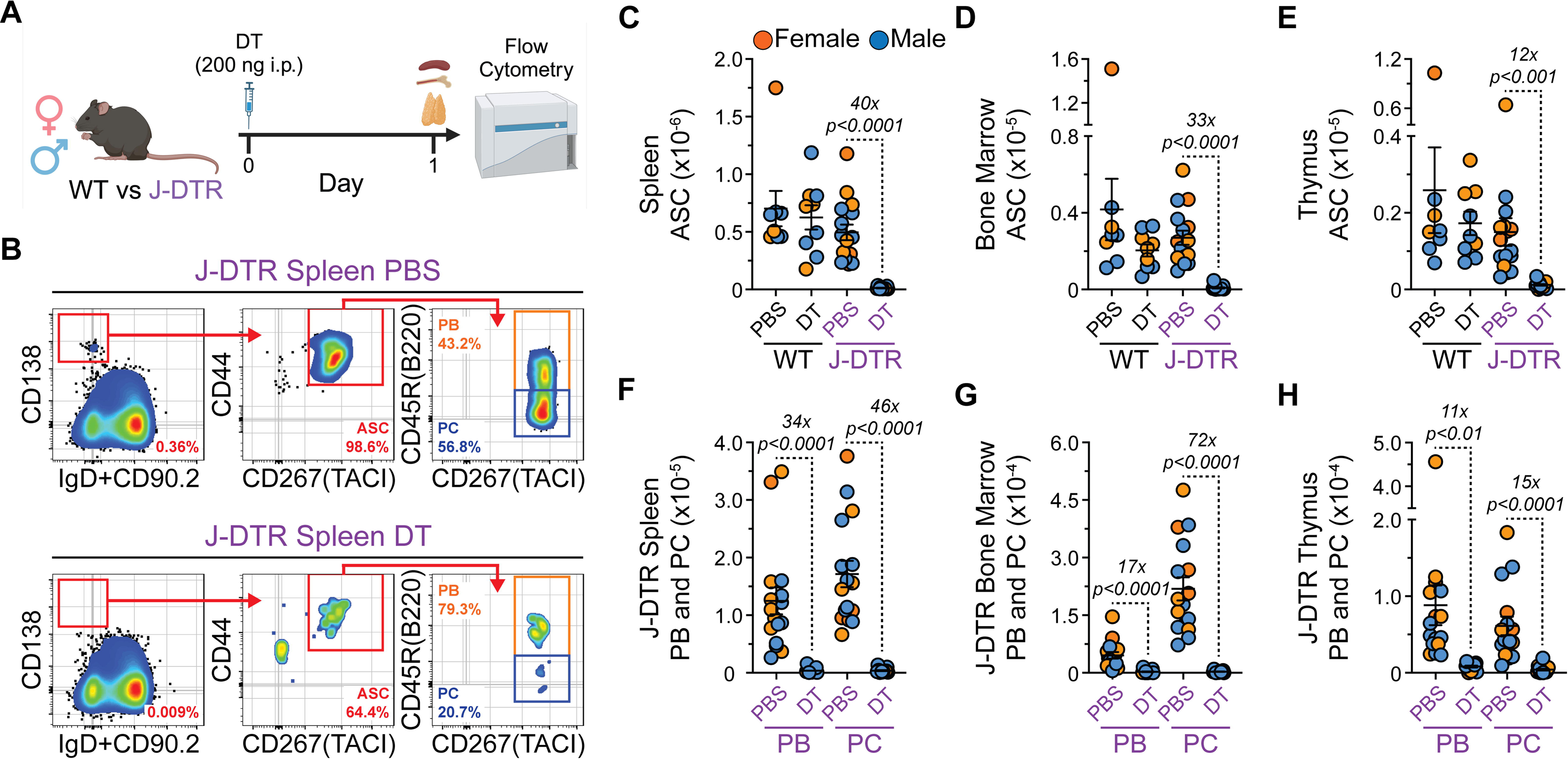
Single dose administration of DT leads to the acute depletion of ASCs in J-DTR mice. **(A)** Schematic showing DT treatment of WT and J-DTR mice. 3-4 months old animals were given a single i.p. dose of 200 ng DT in 100 μL 1x PBS. Control mice received 100 μL of 1x PBS. Mice were euthanized the next day and spleen, bone marrow and thymus were assessed for ASCs and other B cell populations via flow cytometry. Schematic made with BioRender. **(B)** Representative flow cytometry pseudocolor plots showing gating of spleen ASCs from J-DTR mice treated with PBS or DT. Cells were initially gated on live singlets and numbers in plots indicate percentages of gated populations within the immediate parent population. **(C-E)** Total ASC numbers for **(C)** spleen, **(D)** bone marrow and **(E)** thymus of WT and J-DTR mice treated with PBS or DT. **(F-H)** PB and PC numbers for **(F)** spleen, **(G)** bone marrow and **(H)** thymus of J-DTR mice treated with PBS or DT. **(C-H)** Symbols represent individual female (orange) and male (blue) mice. Horizontal lines represent mean ± SEM. **(C-E)** WT PBS: female n = 3, male n = 5; WT DT: female n = 4, male n = 5; J-DTR PBS: female n = 8, male n = 8; J-DTR DT: female n = 8, male n = 9. Statistics: Unpaired Student’s t-test with comparisons made between PBS and DT treatments within a genotype. **(F-H)** J-DTR PBS: female n = 8, male n = 8; J-DTR DT: female n = 8, male n = 9. Statistics: Unpaired Student’s t-test with comparisons made between PBS and DT treatments within an ASC subset.

**Figure 6:**
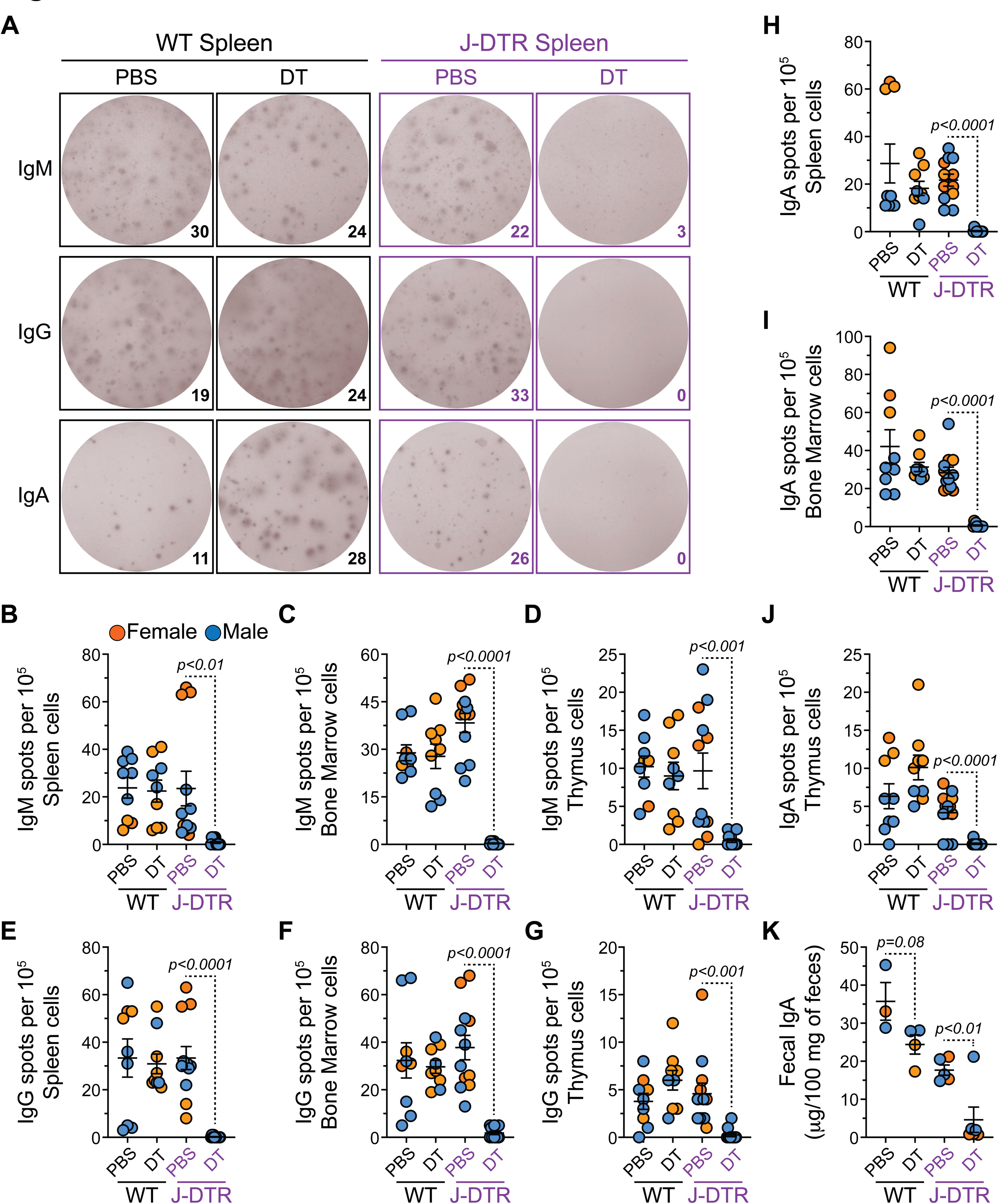
Single dose administration of DT leads to the loss of Ab spot formation and fecal IgA in J-DTR mice. **(A)** Representative ELISpot images showing IgM, IgG and IgA spot formation from spleen of WT and J-DTR mice treated with PBS or DT. 10^5^ cells per well were plated in triplicate. Numbers indicate spots counted per well following background subtraction of identical wells not coated with capture Ab. **(B-D)** Numbers of IgM spots per 10^5^ cells from **(B)** spleen, **(C)** bone marrow and **(D)** thymus of WT and J-DTR mice treated with PBS or DT. **(E-G)** Numbers of IgG spots per 10^5^ cells from **(E)** spleen, **(F)** bone marrow and **(G)** thymus of WT and J-DTR mice treated with PBS or DT. **(H-J)** Numbers of IgA spots per 10^5^ cells from **(H)** spleen, **(I)** bone marrow and **(J)** thymus of WT and J-DTR mice treated with PBS or DT. **(K)** Fecal IgA concentration from WT and J-DTR mice treated with PBS or DT. **(B-J)** Symbols represent individual wells plated in triplicate from 3-5 months old female (orange) and male (blue) WT and J-DTR mice. Values shown represent data following subtraction of spots detected in wells that were not coated with capture Ab (background). Some symbols overlap due to having the same value (e.g., 0). Horizontal lines represent mean ± SEM. WT PBS: female n = 1, male n = 2; WT DT: female n = 2, male n = 1; J-DTR PBS: female n = 2, male n = 2; J-DTR DT: female n = 1, male n = 3. Statistics: Unpaired Student’s t-test with comparisons made between PBS and DT treatments within a genotype. **(K)** Symbols represent individual 3-5 months old female (orange) and male (blue) WT and J-DTR mice. Horizontal lines represent mean ± SEM. WT PBS: female n = 1, male n = 2; WT DT: female n = 2, male n = 2; J-DTR PBS: female n = 3, male n = 2; J-DTR DT: female n = 3, male n = 3. Statistics: Unpaired Student’s t-test with comparisons made between PBS and DT treatments within a genotype.

Phenotyping of J-DTR mice indicated that mIgA^+^ ASCs expressed the highest levels of DTR (**Figures S4C-S4E**). To determine if DT selectively ablated IgA ASCs, we performed mIgM versus mIgA flow cytometry on ASCs from the spleen, bone marrow and thymus of J-DTR mice that received either PBS or DT (**Figures S6K-S6M**). DT injection led to a reduction in mIgM^+^, mIgA^+^ and DN ASCs in the spleen (**Figure S6K**), bone marrow (**Figure S6L**) and thymus (**Figure S6M**). Based upon cell numbers, it appeared that mIgA^+^ ASCs were more efficiently depleted in the spleen (mIgM^+^: 30x fold decrease; mIgA^+^: 773x fold decrease; DN: 285x fold decrease) and bone marrow (mIgM^+^: 20x fold decrease; mIgA^+^: 211x fold decrease; DN: 51x fold decrease) (**Figures S6K-S6L**). In the thymus, mIgM^+^ (16x fold decrease) and mIgA^+^ (12x fold decrease) ASCs were depleted at a relatively similar magnitude with thymus DN ASCs being reduced by ∼7x (**Figures S6M**).

Until now, we have measured depletion based upon flow cytometry and cellular identification via cell surface markers. Thus, we performed ELISpot assays (**Figures 6A-J**) on the spleen, bone marrow and thymus to provide functional confirmation of ASC depletion. In all 3 organs analyzed, administration of DT to J-DTR mice resulted in a significant reduction in the numbers of IgM, IgG and IgA spots per 10^5^ cells (**Figures 6B-6J**). DT treatment did not significantly impact the numbers of spots observed using cells from WT animals (**Figures 6B-6J**). A previous study has demonstrated mouse serum Ab half-lives to range from 17-22 hours (polymeric IgA) to days depending on IgG isotype^33^. Based on those observations, we did not evaluate plasma Abs in our mice as DT treatment lasted less than 24 hours from a practical standpoint. However, we did hypothesize that fecal IgA levels may be reduced in DT-treated J-DTR mice as the concentration of fecal IgA represents a balance between production and excretion. While injection of DT did not significantly alter fecal IgA in WT mice (**Figure 6K**), J-DTR animals that received DT displayed a significant reduction in fecal IgA (**Figure 6K**) most likely also indicative of gut ASC depletion.

Finally, since we previously observed low level DTR expression by selected upstream B cell populations, we also evaluated how DT treatment impacted these populations (**Figures S7A-S7E**). Overall, spleen and thymus total B cells were not impacted by DT treatment (**Figures S7B-S7C**). However, J-DTR mice demonstrated a reduction in their spleen GCB numbers following administration of DT (∼8.8x, **Figure S7D**) supporting the above observation of DTR expression in at least a portion of spleen GCBs. While this level of depletion was significant, it was far less than that observed for spleen ASCs (∼40x, **Figure 5C**) from the same animals. In contrast, DT had no impact on thymus GCB-like cells (**Figure S7E**). Taken together, these data demonstrate that DT can induce acute ASC depletion in J-DTR mice with a limited impact on other mature B cells populations especially in the thymus.

### DT treatment of J-DTR mice allows for kinetic analysis of ASC production

Understanding the kinetics of ASC formation and how newly generated ASCs compete with pre-existing cells for survival niches are important considerations in vaccine development. To determine if our J-DTR mice provide a suitable model to evaluate ASC differentiation kinetics, we treated 3-4 months old female and male J-DTR mice with either PBS or DT (200 ng) (**Figure 7A** and **Figure S8A**). Subsequently, animals were euthanized at days 1, 3 and 7 post-treatment with B cell and ASC populations from spleen, bone marrow and thymus being analyzed by flow cytometry (**Figure 7A** and **Figure S8A**). Similar to above, DT treatment did not alter spleen, bone marrow or thymus cellularity in J-DTR mice (**Figures 7B-7D**)

**Figure 7:**
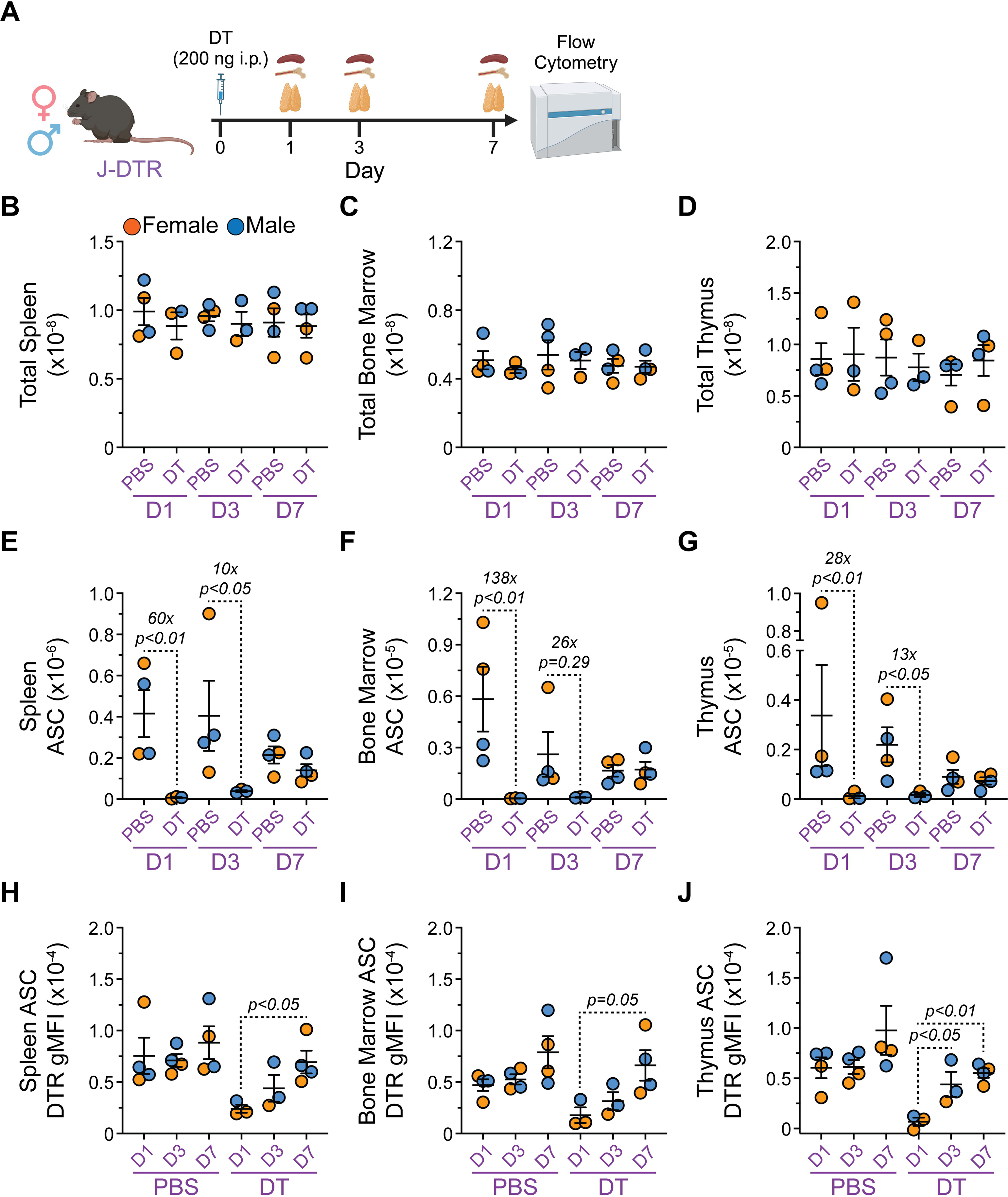
Single dose DT administration allows for the assessment of ASC reconstitution kinetics in J-DTR mice. **(A)** Schematic showing DT treatment of J-DTR mice. 3-4 months old animals were given a single i.p. dose of 200 ng DT in 100 μL 1x PBS. Control mice received 100 μL of 1x PBS. Mice were euthanized at days 1, 3 and 7 post-injection. Spleen, bone marrow and thymus were assessed for ASCs via flow cytometry. Schematic made with BioRender. **(B-D)** Total cell numbers for **(B)** spleen, **(C)** bone marrow and **(D)** thymus of J-DTR mice treated with PBS or DT. **(E-G)** Total ASC numbers for **(E)** spleen, **(F)** bone marrow and **(G)** thymus of J-DTR mice treated with PBS or DT. **(H-J)** DTR gMFIs for ASCs from **(H)** spleen,, **(I)** bone marrow and **(J)** thymus of J-DTR mice treated with PBS or DT. **(B-J)** Symbols represent individual female (orange) and male (blue) mice. Horizontal lines represent mean ± SEM. J-DTR day 1 PBS: female n = 2, male n = 2; J-DTR day 1 DT: female n = 2, male n = 1; J-DTR day 3 PBS: female n = 2, male n = 2; J-DTR day 3 DT: female n = 1, male n = 2; J-DTR day 7 PBS: female n = 2, male n = 2; J-DTR day 7 DT: female n = 2, male n = 2. **(B-G)** Statistics: Kruskal-Wallis test (nonparametric) with Dunn’s multiple comparisons test. Comparisons made between PBS and DT treatments for a given day. **(H-J)** Statistics: One-way ANOVA with Tukey’s multiple comparisons test. Comparisons made between D1, D3 and D7 for a given treatment.

Consistent with previous data (**Figure 5**), DT significantly reduced spleen ASCs in J-DTR mice at 1-day post-injection (∼60x, **Figure 7E**) with ASCs still being depleted ∼10x at day 3 (**Figure 7E**). By day 7, spleen ASC numbers resembled those in PBS-treated animals (**Figure 7E**). Within the bone marrow, ASCs were acutely depleted ∼138x at day 1 and ultimately reached levels observed in control mice by day 7 (**Figure 7F**). At day 3, bone marrow ASCs appeared to still be reduced in DT-treated animals although this did not reach statistical significance (**Figure 7F**). Finally, administration of DT resulted in reduced thymus ASC numbers at both day 1 (∼28x, **Figure 7G**) and day 3 (∼13x, **Figure 7G**). Similar to the spleen and bone marrow, thymus ASCs returned to normal levels by day 7 post-DT (**Figure 7G**). Interestingly enough, ASCs from DT-treated J-DTR mice expressed lower DTR at day 1 compared to day 7 post-treatment in all 3 organs assessed (**Figures 7H-7J**). As with the acute depletion studies (**Figure S7**), we also assessed the effects of DT-treatment on total B cell and GCB (or GCB-like) populations in the spleen and thymus of J-DTR animals (**Figures S8A-S8E**). The only notable changes were reflected in a significant reduction in J-DTR spleen GCBs at 1 day following DT administration (**Figure S8D**) which was followed by a non-significant reduction at day 3. Spleen GCBs were at normal levels at day 7 post-DT treatment (**Figure S8D**). Collectively, these data demonstrate that ASCs are continuously replenished in the spleen, bone marrow and thymus of young mice and that our J-DTR mouse model provides a suitable platform to study ASC reconstitution kinetics.

## Discussion

The data presented here outline the creation and validation of a mouse model in which the endogenous *Jchain* locus drives *DTR* gene expression (J-DTR mice). As shown, ASCs from J-DTR mice express high amounts of DTR protein on the cell surface and can be acutely depleted following a single dose of DT. Furthermore, due to the short half-life of DT, we were able to demonstrate that these mice provide a platform to assay ASC differentiation kinetics following their initial ablation. Finally, we performed all experiments using both sexes showing that this model could be utilized to study ASCs in both females and males.

In generating a mouse strain to allow for the acute deletion of ASCs, we ultimately chose to insert the primate *DTR* coding sequence into the endogenous mouse *Jchain* locus while maintaining the fidelity of exons required for full-length *Jchain* expression. This genomic location was suitable in part due to the high expression of *Jchain* in ASCs relative to other immune cells (**Figure S1**) and^11^. In addition, data seem to indicate that *Jchain* is highly expressed in all ASC subsets, although magnitude differences exist between Ab isotypes^29^. Along these lines, we observed the highest DTR expression (i.e., gMFI) by mIgA^+^ ASCs (**Figure S4**). Jchain plays a pivotal role in the production of pentameric IgM and dimeric IgA and genetic ablation of *Jchain* leads to significant alterations in IgA and to a lesser extent IgM^31,32,34-36^. While there has been some phenotypic variation between previous *Jchain*^-/-^ models, the most reproducible phenotypes consisted of increased circulating IgA and decreased IgA secretion into the gut lumen. The latter of which most likely being due to an inability of dimeric IgA to form and associate with the secretory chain^31,36^ rather than a loss of IgA producing ASCs in the gut^32,35^. Although no developmental defects in J-DTR ASC production in the spleen, bone marrow or thymus were observed (**Figure 4**), we noted an ∼4.5x increase in plasma IgA (**Figure 4J**) and a concomitant ∼2x decrease in fecal IgA (**Figure 4K**) in J-DTR animals. This result was surprising as analysis of ASC-enriched samples from J-DTR spleens for both *Jchain* mRNA (**Figure 2G**) and protein (**Figure S3B**) displayed no obvious deficiencies relative to WT. However, we did not directly assay gut ASCs for *Jchain* levels due to technical limitations. All that being said, the IgA phenotypes present in our J-DTR mice are quite mild compared to those observed in *Jchain*^-/-^ mice^32,35,36^ possibly due to our only using heterozygotes for the J-DTR allele in this study. It will be interesting to examine J-DTR homozygous mice in future studies to determine if differences in IgA localization are further exaggerated which may indicate altered Jchain protein function because of the J-DTR allele. In addition, experiments using cholera toxin vaccination to induce a gut IgA response followed by challenge with *Vibrio cholerae*^37^ may shed light as to the functional consequences resulting from altered IgA distribution in J-DTR mice.

It is difficult to compare models without testing them side-by-side. However, our J-DTR mice appear to be at least as effective as the recently published BICREAD^7^ and CD138-DTR^15^ mouse strains. This was demonstrated by the ability to deplete ASCs within 1 day following a single injection of DT. Although, we did not reach 100% depletion following a single dose of DT which may be a result of the previously reported limited DT half-life^19^ combined with the time between DT treatment and terminal harvest which was ∼15-18 hours for our 1 day depletion studies. It is possible that repetitive DT injections would have better “saturated the system” allowing for more complete depletion. Regardless, ASC depletion was still highly significant and clearly surpassed in magnitude what was previously achieved with antibody-mediated targeting of ASCs^12^. Critically, we observed potent depletion of all ASC isotypes examined in the spleen, bone marrow and thymus which included those expressing IgM, IgG and IgA Abs (**Figure 6** and **Figure S6**). These results correlated well with previously published studies that performed tamoxifen-induced tandem-dimer red fluorescent protein (tdRFP) fluorescent labelling of ASC subsets using *Jchain*-CreERT2 animals^11^. Furthermore, we used fecal IgA concentration as a proxy for gut ASCs and observed a significant reduction in all but 1 J-DTR mouse which received DT (**Figure 6K**). These data support the ability of intraperitoneal administration of DT to eliminate ASCs in a wide range of tissues bolstering the technical and experimental utility of the J-DTR mouse strain.

To demonstrate the feasibility of this model in terms of studying ASC differentiation kinetics, we administered a single dose of DT to J-DTR mice and assayed ASC numbers 1-, 3- and 7-days post-treatment (**Figure 7**). While these experiments were limited in power, they clearly showed the ability to observe ASC reconstitution partially in the spleen at 3 days post-DT with ASC numbers returning to normal by 7 days. Reconstitution in the bone marrow and thymus was also complete by 7 days. As B cell activation and ASC production in the thymus is particularly relevant to myasthenia gravis (MG)^38,39^, the ability to deplete thymus ASCs in J-DTR mice and study their generation within the organ may provide a critical tool that can be used to understand MG etiology^40^. Furthermore, we previously demonstrated that thymus ASCs possessed major histocompatibility complex class II (MHC II) on their cell surface and transcriptionally expressed machinery required for antigen presentation^27^. Hence, using this model to deplete thymus ASCs long-term or over discrete developmental windows may prove informative regarding their potential to regulate T cell development and selection in the thymus.

An interesting observation was that J-DTR ASCs remaining 1 day following DT treatment expressed low to intermediate levels of surface DTR compared to ASCs from PBS-treated J-DTR mice (**Figure S6**). While it is possible that these cells would never express high DTR levels, we suspect that their low expression may have been a sign of overall immaturity as the ASCs present after the administration of DT had increased PB:PC ratios (**Figure S6**). In alignment with this, we observed the lowest DTR levels by ASCs from J-DTR mice immediately following DT treatment when compared to those present 7 days post-DT (**Figure 7**). However, this is purely speculative and integration of a reporter strain allowing for genetic timestamping would enable us to definitively address the above results. In this sense, ASCs would be fluorescently timestamped prior to depletion. If fluorescently labeled ASCs are still present following DT treatment and lack DTR, then this may be indicative of a rare, minority population not expressing high amounts of *Jchain* and thus lacking sufficient amounts of DTR required for ablation. In contrast, if ASCs lacking DTR are not labelled, then these cells could represent nascently generated ASCs which would be examined for maturation status using metrics such as Prdm1 expression levels which increases with maturity^28,41^ and Ki-67 which would be lost as PBs transition to a more mature PC phenotype^27,28^.

While no single model is perfect, the J-DTR model provides an alternative platform to deplete ASCs acutely. This is particularly relevant as DT treatment of BICREAD mice would presumably also target *Prdm1* expressing tissue resident and/or memory T cell subsets^16-18^ while CD138-DTR mice may possess experimental caveats due to potentially targeting a subset of IL-10 producing CD138^+^ macrophages^42^. Administration of DT to J-DTR mice did not result in alterations in organ cellularity of the spleen, bone marrow and thymus or even total B cell populations in the spleen and thymus. This was an important result and indicated that widespread leaky expression of DTR was not present. However, we did see a reduction in J-DTR spleen GCBs upon DT treatment which may have been a direct effect as we were able to detect low levels of surface DTR expression on these cells. This observation is consistent with recent data using a *Jchain*-driven CreERT2 in combination with a tandem-dimer Tomato (tdTomato) reporter that labelled GCBs following West Nile virus vaccination^43^. Similarly, the use of a mouse strain with an enhanced green fluorescent protein (eGFP) reporter and CreERT2 inserted into the endogenous *Jchain* locus demonstrated the dominant presence of GCBs within the eGFP^LO^ B220^HI^ cell population following sheep red blood cell immunization^11^. In this instance, labelling with tdRFP (i.e., Cre activity) following tamoxifen induction^11^ did not appear as penetrant compared to that observed in the West Nile virus study^43^. Multiple variables may account for these outcomes such as antigen type and load, tamoxifen dosage as well as differences in tdTomato versus tdRFP detection. Relevant to the current study, experiments performed by Xu *et al*.^11^ indicated that GFP positivity was overall quite low in both dark zone and light zone GCBs suggesting that only a limited proportion of these cells express *Jchain*. In contrast to spleen GCBs, GCB-like cells in the thymus did not express appreciable levels of DTR nor were they depleted by DT treatment. In this study, we defined thymus GCB-like cells as expressing both CD95(Fas) and GL7. These cells possess similarities to the GL7^+^ CD38^+^ B cell subset previously identified in the thymus^44^ and may be more akin to an activated memory B cell phenotype^45^. In terms of the spleen, the off-target depletion of GCBs was ∼8.8x (**Figure S7D**) whereas the specific and intended elimination of ASCs was ∼40x (**Figure 5C**) indicating a differential activity of DT treatment as expected. Whether the low-level elimination of GCBs is relevant to the measure of ASC differentiation kinetics following immunization will most likely be context dependent and require the inclusion of a DT-treated J-DTR control group naïve to the immunogen of interest being studied. Ultimately, it will be interesting to see if modulating the dose or treatment regimen of DT (single dose of 200 ng used here) can further exploit the ability of DT to specifically target ASCs over other B cell populations using our J-DTR model. That being said, we have clearly shown that the J-DTR mouse model represents a new genetic platform that can be leveraged to study ASC production, and potentially even function, in a variety of experimental contexts.

## Supporting information

Key Resources Table

## Resource Availability

### Lead Contact

Further information and requests for resources and reagents should be directed to and will be fulfilled by the Lead Contact, Peter Dion Pioli.

### Materials Availability

Materials underlying this article will be shared by the lead contact upon request. The J-DTR mice have not been deposited in an animal repository. However, J-DTR animals will be made available by the Pioli laboratory upon completion of a material transfer agreement.

### Data and Code Availability

- Flow cytometry data reported in this study will be shared by the lead contact upon request.
- Any information required for data reanalysis is available from the lead contact upon request.

## Limitations of the Study

Regarding limitations, the work presented here focused on validating and establishing the J-DTR mouse model as a tool to study ASC differentiation. While our experiments evaluated the functionality of J-DTR mice at homeostasis, it is expected that DTR-expressing ASCs would continue to be sensitive to DT treatment even in the context of infection-based mouse models, although this remains to be tested. Furthermore, additional experiments will be required to demonstrate the efficacy of long-term ASC depletion using repetitive DT administration which will be particularly important as immune responses in the context of repeated DT injections have been previously noted^46^. In this case, combination of the aforementioned *Jchain*-CreERT2 strains with a Flox-STOP-Flox diphtheria toxin fragment A allele^47^ may provide an alternative targeting strategy.

## Acknowledgments

Funding was provided by the University of Saskatchewan College of Medicine via intramural startup funds and the Office of the Vice Dean Research College of Medicine Research Award (CoMRAD). This work was further supported by the National Institute on Aging of the National Institutes of Health under Award Number R03AG071955, Saskatchewan Health Research Foundation (Establishment Grant, Award Number 6230) and the Natural Sciences and Engineering Research Council of Canada (Discovery Grant, Award Number 2024 400 06646). The content is solely the responsibility of the authors and does not necessarily represent the official views of any funding sources. Graphical Abstract (https://BioRender.com/k32p032), Figure 1A (https://BioRender.com/9yo57vc), Figures 5A, S6A and S7A (https://BioRender.com/0r1fj9f) as well as Figures 7A and S8A (https://BioRender.com/k6h9915) were constructed using BioRender.

## Author Contributions

K.T.P. and P.D.P designed experiments. K.T.P., M.R., H.H. and P.D.P. conducted and analyzed experiments.

K.T.P. and P.D.P wrote the manuscript, and all authors approved of the manuscript.

## Supplemental Information

Document S1. Figures S1-S8

## STAR Methods

### EXPERIMENTAL MODEL AND SUBJECT DETAILS

#### Experimental Animals

J-DTR mice were originally generated fee-for-service by InGenious Targeting Laboratory. Upon receipt, animals were quarantined and verified pathogen and parasite free before being released for use. Animals were subsequently bred and maintained at the University of Saskatchewan Lab Animals Services Unit. Wildtype (WT) and *Jchain*^+/*DTR*^ heterozygote female and male mice were utilized for all experiments. Animals ranged in age from 3-7 months when used for experiments. Animal care and use were conducted according to the guidelines of the University of Saskatchewan University Animal Care Committee Animal Research Ethics Board and all experiments were approved under Animal Use Protocol 20220020.

### METHOD DETAILS

#### Creation of J-DTR mice

This mouse model was created using a genetically engineered mouse embryonic stem cell line, in which a custom targeting vector was designed so that the internal ribosomal entry site (IRES)-*DTR* cassette was inserted after the TAG stop codon of the *Jchain* gene. The knock-in cassette was followed by an FRT-flanked neomycin (Neo) selection cassette. The long homology arm (LA) of the vector is ∼6 kb in length and the short homology arm (SA) is ∼2.1 kb in length. The region used to construct the targeting vector was subcloned from a positively identified C57BL/6 BAC clone using homologous recombination-based techniques. The targeting vector was confirmed by restriction analysis and sequencing after each modification step.

The targeting vector was then linearized and electroporated into a C57BL/6 (BF1) embryonic stem cell (ESC) line expressing flippase (FLP) recombinase. After selection with G418, antibiotic-resistant colonies were picked, expanded and screened via PCR analysis and sequenced for homologous recombinant ESC clones. The Neo resistance cassette was removed via FLP recombinase in the ESCs during expansion. Positively targeted ESC clones were then microinjected into BALB/c blastocysts and transferred into pseudo-pregnant females. Resulting chimeras with a high percentage black coat color were mated to C57BL/6N WT mice, after which the offspring were tail-tipped and genotyped for germline transmission of the targeted allele sequence. Germline mice were identified as heterozygous for the co-expression of the *DTR* cassette in the mouse *Jchain* gene locus. Upon receipt, mice were bred to eliminate the gene encoding the FLP.

#### Genomic DNA Isolation and Genotyping

Ear biopsies were incubated at 100°C in 400 μL of 50 mM sodium hydroxide (NaOH) until tissue was fully dissolved. NaOH was neutralized by the addition of 1 M Tris-hydrochloric acid (HCl), pH 8.0 (50 μL). Samples were vortexed and then centrifuged at 25°C and 12,000g for 2 minutes. Supernatant (200 μL) was transferred to a new 1.5-mL tube. After the addition of 3 M sodium acetate (NaOAc), pH 5.2 (20 μL) and 95% ethanol (EtOH) (660 μL), samples were vortexed, and DNA was precipitated overnight (O/N) at −20°C. The next day, samples were centrifuged at 4°C and 12,000g for 5 minutes. Supernatant was aspirated and DNA pellets were resuspended in 100 μL of 0.1x Tris-EDTA buffer.

For genotyping PCR, each reaction consisted of 10 μL of 2x Platinum II Hot-Start PCR Master Mix (Thermo Fisher Scientific, Cat# 14000012), 1 μL forward primer, 1 μL reverse primer, 2 μL DNA and 6 μL H_2_O. All primers were resuspended at a concentration of 1 μg/mL in 0.1x TE. Reactions were amplified using a Veriti 96-well thermal cycler (Thermo Fisher Scientific). Reactions products were electrophoresed in 2% agarose gels containing ethidium bromide and products were visualized under ultraviolet light using a Bio-Rad ChemiDoc Imaging System. All genotyping primer sequences are listed in the **Key Resources Table** and PCR amplification was performed as follows: Stage 1 (1 cycle) - 94°C for 2 minutes; Stage 2 (40 cycles) - 94°C for 30 seconds, 62°C for 30 seconds, 72°C for 1 minute; Stage 3 (1 cycle) - 72°C for 10 minutes. Samples were stored at 4°C until agarose gel electrophoresis.

#### In Vivo DT Treatment

1 mg of lyophilized DT from *Corynebacterium diphtheriae* (Millipore Sigma, Cat# D0564) was resuspended in 0.5 mL sterile H_2_O yielding a 2 mg/mL DT concentration in a 10 mM Tris-1mM EDTA, pH 7.5 solution. For injection, DT was subsequently diluted to 2 μg/mL in 1x phosphate buffered saline (PBS) (Gibco, Cat# 21600-069). Mice received 100 μL intraperitoneal (i.p.) injections of either PBS or DT (200 ng total).

#### Isolation of Bone Marrow, Spleen and Thymus Tissue

All tissues were processed and collected in calcium and magnesium-free 1x PBS. Spleen and thymus were dissected and crushed between the frosted ends of two slides. Bone marrow was isolated from both femurs and tibias by cutting off the end of bones and flushing the marrow from the shafts and ends using a 23-gauge needle. Cell suspensions were centrifuged for 5 minutes at 4°C and 600g. Red blood cells were lysed by resuspending cells in 3 mL of 1x red blood cell lysis buffer on ice for ∼3 minutes. Lysis was stopped with the addition of 7 mL of 1x PBS. Cell suspensions were strained through 40 μm filters and counted on a Countess 3 (Thermo Fisher Scientific) using Trypan Blue to exclude dead cells. Cell suspensions were centrifuged as before (5 minutes at 4°C and 600g) and resuspended at 2×10^7^ cells/mL in 1x PBS + 0.1% bovine serum albumin (BSA, Fisher BioReagents, Cat# BP9706-100) before use.

#### Isolation of Fecal Pellet Supernatants

Fecal pellets were collected from dissected mouse intestines and colon. Tools were used to open the organs and collect fecal pellets into 1.5-mL microtubes. The collected pellets were placed on ice until generation of fecal pellet suspensions. To generate suspensions, pellets were weighed and 1x PBS was added at 1 mL per 100 mg of feces. Tubes were vortexed at max speed in a 4°C cold room for 10 minutes. The fecal suspensions were centrifuged at 12,000g for 10 min at 4°C. Then the supernatant was transferred into a new microtube and stored at −80°C until use.

#### Isolation of Plasma

Blood was initially collected into 1.5-mL microtubes via cardiac puncture then placed on ice until completion of tissue harvesting. Subsequently, blood was set at room temperature (RT) for 30 min and then centrifuged at 1000g for 10 min in 4°C. Plasma was transferred into a new microtube and stored at −80°C until use.

#### B cell and ASC Enrichment

EasySep Mouse Pan-B cell Isolation and EasySep Release Mouse CD138 Positive Selection kits from STEMCELL Technologies were used to enrich B cells and ASCs from ∼5×10^7^ spleen cells following manufacturer guidelines. Isolated cells were collected in a final volume of 1.5 mL of 1x PBS + 2% fetal bovine serum + 1 mM EDTA and counted using a Countess 3 with Trypan Blue to exclude dead cells and calculate final yield.

#### Quantitative Polymerase Chain Reaction (qPCR)

RNA was extracted from isolated B cells and ASCs using the PureLink RNA Mini Kit (Thermo Fisher Scientific, Cat# 12183025). RNA was quantified using a NanoDrop One^c^ (Thermo Fisher Scientific) and verified to have a A260/280 ratio of ∼2.0. The Maxima H-Minus First Strand cDNA Synthesis Kit with dsDNAse (Thermo Fisher Scientific, Cat# K1682) was used to generate cDNA. Each cDNA synthesis reaction included ≥10 ng RNA mixed with 1 μL random hexamers primers, 1 μL 10 mM dNTP mix, 4 μL RT buffer, 1 μL Maxima H-minus enzyme mix and the appropriate volume of water required to obtain a 20 μL reaction. The reaction mixtures were incubated in a Veriti 96-well thermal cycler using the manufacturer recommended amplification program. qPCR was performed using a StepOnePlus Real-Time PCR System (Applied Biosystems). Each 20 μL reaction contained 2x TAQMAN Fast Advanced Master Mix (10 μL), 20x TaqMan primer (1 μL), cDNA (2 μL) and water (7 μL). Triplicate reactions were run in 96-well plates using standard TAQMAN amplification conditions. All primers are listed in the **Key Resources Table**. Expression of target genes in **Figures 2C-2E** was calculated as 2^(Actb _CT_ – Target_CT_)^ and represents the average derived from triplicate technical replicates. In **Figures 2F-2G**, relative expression values of *DTR* or *Jchain* were divided by those of *Prdm1* to normalize for ASC content in the indicated samples.

#### Immunostaining

All staining procedures were performed in 1x PBS + 0.1% BSA. Samples were labeled with a CD16/32 Ab to eliminate non-specific binding of Abs to cells via Fc receptors. All Abs utilized are listed in the **Key Resources Table**. Cells were incubated on ice for 30 minutes in the dark with the appropriate Abs. Unbound Abs were washed from cells with 1x PBS + 0.1% BSA followed by centrifugation for 5 minutes at 4°C and 600g. Supernatants were decanted, and cell pellets were resuspended in an appropriate volume of 1x PBS + 0.4% BSA + 2 mM EDTA for flow cytometric analysis. Before analysis, cells were strained through a 40 μm filter mesh and kept on ice in the dark. eBioscience Fixable Viability (Live-Dead) Dye eFluor 780 (Thermo Fisher Scientific, Cat# 65-0865-14) was added to samples to assess dead cell content. The stock solution was diluted 1:250 in 1x PBS and 10 μL was added to ∼5 x 10^6^ cells per stain. Live-Dead stain was added concurrent with surface staining Abs.

#### Flow Cytometry

Flow cytometry was performed on a CytoFLEX (Beckman Coulter) located in the Cancer Cluster at the University of Saskatchewan. Total cells were gated using side scatter area (SSC-A) versus forward scatter area (FSC-A). Singlets were identified using sequential gating of forward scatter-height (FSC-H) versus FSC-A and side scatter-height (SSC-H)versus SSC-A. Live cells were Live-Dead stain negative. All data were analyzed using FlowJo (v10) software.

#### Enzyme-Linked Immunosorbent Spot (ELISpot)

ELISpot plates (Millipore Sigma, Cat# MSIPS4W10) were briefly incubated at RT for 1 minute with 15 μL of 35% EtOH. EtOH was removed and wells were washed 3 times with 150 μL of 1x PBS. Subsequently, wells were coated O/N at 4°C with 100 μL of capture Ab. The capture Ab (Millipore Sigma, Cat# SAB3701043-2MG) recognized mouse total IgG+IgM+IgA isotypes and was pre-diluted in 1x PBS to a final concentration of 5 μg/mL before use. The next day, coating Abs were removed, and wells were washed 3 times with 150 μL RPMI 1640. Plates were subsequently blocked with 150 μL RPMI for a minimum of 2 hours at 37°C in a 5% CO_2_/20% O_2_ tissue culture incubator. Blocking solution was removed and total cells from bone marrow, spleen and thymus were loaded into wells with a target number of 10^5^ cells per well in a 100 μL volume (3 wells per sample). Cells had previously been resuspended at 1×10^6^ cells/mL in RPMI supplemented with a proliferation-inducing ligand (APRIL) (10 ng/mL), interleukin (IL)-6 (10 ng/mL), heat-inactivated fetal calf serum (10%), Penicillin-Streptomycin (100 U/mL), L-glutamine (2 mM), Gentamicin (50 μg/mL), sodium pyruvate (1 mM), non-essential amino acids (1x), non-essential vitamins (1x) and 2-mercaptoethanol (10^-5^ M). Cells were then incubated O/N (>12 hours) at 37°C in a 5% CO_2_/20% O_2_ tissue culture incubator. The next day, culture supernatants and cells were removed. Wells were washed 3 times with 150 μL of 1x PBS then an additional 3 times with 150 μL of 1x PBS + 0.1% Tween-20 + 1% BSA. Secondary Abs conjugated to horseradish peroxidase (HRP) were added at a volume of 100 μL per well and plates were incubated for 2 hours at RT. Anti-IgM-HRP (Thermo Fisher Scientific, Cat# 62-6820) and anti-IgA-HRP (Thermo Fisher Scientific, Cat# 62-6720) were both diluted at 1:1000, and anti-IgG-HRP (SouthernBiotech, Cat# 1015-05) was diluted at 1:50,000. All secondary Abs were diluted in 1x PBS + 0.1% Tween-20 + 1% BSA before use. Following incubation, Abs were removed, and plates were washed 3 times with 150 μL of 1x PBS + 0.1% Tween-20 + 1% BSA followed by an additional 3 washes with 150 μL of 1x PBS. To reveal “spots”, 100 μL of Developing Solution from the AEC Substrate Set (BD Biosciences, Cat# 551951) was added to each well. Plates were shaken at 200 rpm for 30 minutes at RT. Developing Solution was removed and plates were washed 5 times with 150 μL of H_2_O. Well backings were removed, and plates dried at RT O/N after which “spots” were visualized with a Mabtech ASTOR ELISPOT reader. Additional wells lacking capture Abs were developed to gauge background. For spot quantification, counting was restricted to a 1250-pixel area of interest to avoid edge artifacts. For each genotype and treatment combination, 3 background wells were counted, averaged and then subtracted to obtain the reported spot numbers.

#### Enzyme-Linked Immunosorbent Assay (ELISA) for Plasma and Fecal Pellet Supernatants

High binding ELISA plates (Greiner, Cat# 655081) were coated using 100 μL anti-mouse IgG/IgA/IgM (H+L) (Sigma Aldrich, Cat# SAB3701043-2MG) at 5 μg/mL in 1x PBS per well and incubated at 4°C O/N covered with plastic wrap. Coating Abs were removed, and plates were washed 3 times with 150 μL 1x PBS + 0.1% Tween 20 (Wash Solution) per well. Subsequently, 150 μL of 1x PBS + 1% BSA + 0.1% Tween (Block Solution) was added per well and plates were blocked at RT for 2 hours. Block Solution was removed, and wells washed 3 times with 150 μL of Wash Solution. Purified Ab standards (Standard Curves) and plasma or fecal supernatant samples were diluted at various concentrations in 1x PBS and 100 μL per dilution was added per well. Plates were incubated at RT for 2 hours; samples/standards were removed and wells washed 3 times with 150 μL of Wash Solution. Isotype-specific HRP-conjugated secondary Abs were diluted in Block Solution and 100 μL were used per well. Secondary Abs for IgM and IgA were diluted to a final concentration of 1:5000 while those for IgG were diluted to 1:50,000. Following a 2-hour RT incubation, wells were washed 3 times with 150 μL of Wash Solution, then incubated for 4 minutes with 100 μL 1x TMB Substrate (Thermo Fisher Scientific, Cat# 00-4201-56) per well. Enzymatic reactions were stopped with the addition of 100 μL per well of 0.16M H_2_SO_4_ (Fisher Chemical, Cat# SA431-500). Optical densities (ODs) were read at 450 nm wavelength using a BioLegend Mini ELISA Plate Reader (BioLegend, Cat# 423555). Samples and standards were analyzed following subtraction of blank wells (1x PBS) and assayed in triplicate. Plasma and fecal supernatant Ab concentrations were calculated using linear portions of standard curves and the equation of a straight line (y = mx + b) where y = average OD per sample, m = slope, x = antibody concentration and b = y-axis intercept. Only experiments with a linear standard curve R^2^ > 0.98 were considered valid.

#### Western Blotting

ASC-enriched samples were generated as indicated in Section 2.8. After counting, cells were centrifuged then lysed using the Thermo Fisher Scientific Cell Extraction Buffer (Cat# FNN0011) following the provided protocol. The only protocol exception was that cells were concentrated 10x in terms of lysis buffer volume. Lysates were stored at −80°C until use. For western blot analysis, sample lysates were combined with Bolt Sample Reducing Agent (10x) (Thermo Fisher Scientific, Cat# B00040) and Bolt LDS Sample Buffer (4x) (Thermo Fisher Scientific, Cat# B0008) then loaded into a Bolt Bis-Tris Mini Protein Gel, 4-12% following the manufacturer’s protocol (Thermo Fisher Scientific, Cat# NW04122BOX). The PageRuler Plus Prestained Protein Ladder (Thermo Fisher Scientific, Cat# 26619) was used as a size standard. Following the Thermo Fisher Scientific Mini Blot Module electrophoresis and blotting protocol, the gel was electrophoresed, and proteins transferred to a PVDF membrane using Invitrogen PVDF/Filter Paper Sandwiches 0.45 μm (Cat# LC2005). After transfer, the PVDF membrane was washed with nuclease-free H_2_O (Thermo Fisher Scientific, Cat# AM9906) and transferred proteins were revealed following staining with 0.1% Ponceau S/5% Acetic Acid for 15 minutes. A cell phone camera image was taken of the Ponceau stained membrane protein. The membrane was washed with Milli-Q H_2_O 3x at 5 minutes per wash then blocked with 1x PBS + 1% Tween + 5% nonfat dry milk (Membrane Blocking Solution) for 1 hour at RT. The membrane was cut above the 35 kDa marker as Jchain possesses a predicted molecular weight of 18 kDa and the bottom third of the membrane was incubated O/N at 4°C with a rabbit anti-human/mouse/rat IgJ (Jchain) (Thermo Fisher Scientific, Cat# 13688-1-AP) primary Ab diluted 1:250 in Membrane Blocking Solution. The following day, the membrane was washed 3x with 1x PBS + 1% Tween before application of a goat anti-rabbit IgG-HRP secondary Ab (Southern Biotech, Cat# 4030-05) diluted 1:10,000 in Membrane Blocking Solution. Following a 2-hour 400 revolutions per minute shaker incubation at RT, the membrane was washed 3x with 1x PBS + 1% Tween. The membrane was developed using a freshly mixed Novex ECL chemiluminescent substrate reagent kit (Thermo Fisher Scientific, Cat#WP20005) for 1 minute before being imaged with a Bio-Rad ChemiDoc using the Chemiluminescent program.

#### Quantification and Statistical Analysis

The numbers of mice used (n =) per experiment are listed in the Figure Legends. Statistical analyses were performed using GraphPad Prism (v8.4.2) software. Quantification of cell numbers and various flow cytometry data are graphically represented as mean ± standard error of the mean (SEM). Statistical analyses are described within each Figure Legend. Unless otherwise stated, only statistically significant p-values (e.g., *p≤0*.*05*) are shown within each Figure.

