## Supplementary material for "Jchain-Diphtheria Toxin Receptor Mice Allow for Diphtheria Toxin-Mediated Depletion of Antibody-Secreting Cells and Analysis of Differentiation Kinetics": Key Resources Table

The table highlights the genetically modified organisms and strains, cell lines, reagents, software, and source data **essential** to reproduce results presented in the manuscript. Depending on the nature of the study, this may include standard laboratory materials (i.e., food chow for metabolism studies), but the Table is **not** meant to be comprehensive list of all materials and resources used (e.g., essential chemicals such as SDS, sucrose, or standard culture media don’t need to be listed in the Table). **Items in the Table must also be reported in the Method Details section within the context of their use.** The number of **primers and RNA sequences** that may be listed in the Table is restricted to no more than ten each. If there are more than ten primers or RNA sequences to report, please provide this information as a supplementary document and reference this file (e.g., See Table S1 for XX) in the Key Resources Table.

***Please note that ALL references cited in the Key Resources Table must be included in the References list.*** Please report the information as follows:

- **REAGENT or RESOURCE:** Provide full descriptive name of the item so that it can be identified and linked with its description in the manuscript (e.g., provide version number for software, host source for antibody, strain name). In the Experimental Models section, please include all models used in the paper and describe each line/strain as: model organism: name used for strain/line in paper: genotype. (i.e., Mouse: OXTR^fl/fl^: B6.129(SJL)-Oxtr^tm1.1Wsy/J^). In the Biological Samples section, please list all samples obtained from commercial sources or biological repositories. Please note that software mentioned in the Methods Details or Data and Software Availability section needs to be also included in the table. See the sample Table at the end of this document for examples of how to report reagents.
- **SOURCE:** Report the company, manufacturer, or individual that provided the item or where the item can obtained (e.g., stock center or repository). For materials distributed by Addgene, please cite the article describing the plasmid and include “Addgene” as part of the identifier. If an item is from another lab, please include the name of the principal investigator and a citation if it has been previously published. If the material is being reported for the first time in the current paper, please indicate as “this paper.” For software, please provide the company name if it is commercially available or cite the paper in which it has been initially described.
- **IDENTIFIER:** Include catalog numbers (entered in the column as “Cat#” followed by the number, e.g., Cat#3879S). Where available, please include unique entities such as [RRIDs](https://www.force11.org/group/resource-identification-initiative), Model Organism Database numbers, accession numbers, and PDB or CAS IDs. For antibodies, if applicable and available, please also include the lot number or clone identity. For software or data resources, please include the URL where the resource can be downloaded. Please ensure accuracy of the identifiers, as they are essential for generation of hyperlinks to external sources when available. Please see the Elsevier [list of Data Repositories](https://www.elsevier.com/authors/author-resources/research-data/data-base-linking) with automated bidirectional linking for details. When listing more than one identifier for the same item, use semicolons to separate them (e.g. Cat#3879S; RRID: AB_2255011). If an identifier is not available, please enter “N/A” in the column.
  - ***A NOTE ABOUT RRIDs:*** We highly recommend using RRIDs as the identifier (in particular for antibodies and organisms, but also for software tools and databases). For more details on how to obtain or generate an RRID for existing or newly generated resources, please [visit the RII](https://www.force11.org/group/resource-identification-initiative) or [search for RRIDs](https://scicrunch.org/resources).

Please use the empty table that follows to organize the information in the sections defined by the subheading, skipping sections not relevant to your study. Please do not add subheadings. To add a row, place the cursor at the end of the row above where you would like to add the row, just outside the right border of the table. Then press the ENTER key to add the row. Please delete empty rows. Each entry must be on a separate row; do not list multiple items in a single table cell. Please see the sample table at the end of this document for examples of how reagents should be cited.

***TABLE FOR AUTHOR TO COMPLETE***

*Please upload the completed table as a separate document.* ***Please do not add subheadings to the Key Resources Table.*** *If you wish to make an entry that does not fall into one of the subheadings below, please contact your handling editor. (****NOTE:*** *For authors publishing in Current Biology, please note that references within the KRT should be in numbered style, rather than Harvard.)*

**KEY RESOURCES TABLE**

| **REAGENT or RESOURCE** | **SOURCE** | **IDENTIFIER** |
| --- | --- | --- |
| **Antibodies** | | |
| CD138-BV421 (Clone: 281-2) | BD Biosciences | Cat# 562610; RRID: AB_11153126 |
| IgD-BV605 (Clone: 11-26c.2a) | BioLegend | Cat# 405727; RRID: AB_2562887 |
| CD90.2(Thy-1.2)-BV605 (Clone: 53-2.1) | BD Biosciences | Cat# 563008; RRID: AB_2665477 |
| CD45R(B220)-BV711 (Clone: RA3-6B2) | BD Biosciences | Cat# 563892; RRID: AB_2738470 |
| CD45R(B220)-PerCP/Cy5.5 (Clone: RA3-6B2) | BD Biosciences | Cat# 552771; RRID: AB_394457 |
| CD19-eFluor 506 (Clone: eBio1D3) | Thermo Fisher Scientific | Cat# 69-0193-82; RRID: AB_2637306 |
| CD16/32-Unlabeled (Clone: 93) | Thermo Fisher Scientific | Cat# 14-0161-86; RRID: AB_467135 |
| CD44-FITC (Clone: IM7) | Thermo Fisher Scientific | Cat# 11-0441-82; RRID: AB_465045 |
| CD44-BV711 (Clone: IM7) | Thermo Fisher Scientific | Cat# 407-0441-82; RRID: AB_2937199 |
| CD267(TACI)-PE (Clone: 8F10) | BioLegend | Cat# 133403; RRID: AB_2203542 |
| GL7-AF647 (Clone: GL7) | BD Biosciences | Cat# 561529; RRID: AB_10716056 |
| CD95(Fas)-PE (Clone: Jo2) | BD Biosciences | Cat# 554258; RRID: AB_395330 |
| IgM-FITC (Clone: polyclonal) | SouthernBiotech | Cat#: 1021-02; RRID: AB_2794237 |
| IgA-APC (Clone: mA-6E1) | Thermo Fisher Scientific | Cat# 17-4204-82; RRID: AB_2848294 |
| DTR-Biotin (Clone: polyclonal) | R&D Systems | Cat# BAF259; RRID: AB_2114598 |
| Streptavidin-PE/Cy7 | BD Biosciences | Cat# 557598 |
| Anti-mouse IgG+IgA+IgM (H+L) | Millipore Sigma | Cat# SAB3701043-2MG |
| Mouse IgM Isotype Control (Clone: 11E10) | Thermo Fisher Scientific | Cat# 14-4752-82; RRID: AB_470123 |
| Mouse IgG Isotype Control (Clone: polyclonal) | Thermo Fisher Scientific | Cat# 02-6502; RRID: AB_2532951 |
| Mouse IgA Isotype Control (Clone: S107) | Thermo Fisher Scientific | Cat# 14-4762-81 RRID: AB_470125 |
| Goat anti-mouse IgM-HRP (Clone: polyclonal) | Thermo Fisher Scientific | Cat# 62-6820;  RRID: AB_2533954 |
| Goat anti-mouse IgG-HRP (Clone: polyclonal) | SouthernBiotech | Cat# 1015-05; RRID: AB_2794194 |
| Goat anti-mouse IgA-HRP (Clone: polyclonal) | Thermo Fisher Scientific | Cat# 62-6720;  RRID: AB_2533951 |
| Rabbit anti-human/mouse/rat IgJ (Jchain) (Clone: polyclonal) | Thermo Fisher Scientific | Cat# 13688-1-AP;  RRID: AB_2877973 |
| Goat anti-rabbit IgG-HRP (Clone: polyclonal) | SouthernBiotech | Cat# 4030-05; RRID: AB_2687483 |
| **Chemicals, Peptides, and Recombinant Proteins** | | |
| Bovine Serum Albumin (DNase- and Protease-free) | Fisher BioReagents | Cat# BP9706-100 |
| Nuclease-free H_2_O | Thermo Fisher Scientific | Cat# AM9906 |
| APRIL | Fisher Scientific | Cat# 7907-AP-010CF |
| IL-6 | Fisher Scientific | Cat# PMC0066 |
| Ponceau S | Fisher BioReagents | Cat# BP103-10 |
| Glacial Acetic Acid | Fisher Chemical | Cat# A38SI-212 |
| Blotting-Grade Blocker (nonfat dry milk) | Bio-Rad | Cat# 1706404 |
| Bolt LDS Sample Buffer (4x) | Thermo Fisher Scientific | Cat# B0008 |
| Bolt Sample Reducing Agent (10x) | Thermo Fisher Scientific | Cat# B0004 |
| NuPAGE MOPS SDS Running Buffer (20x) | Thermo Fisher Scientific | Cat# NP0001 |
| Bolt Transfer Buffer (20x) | Thermo Fisher Scientific | Cat# BT0006 |
| Halt Protease & Phosphatase Inhibitor  Single-use Cocktail (100X) | Thermo Fisher Scientific | Cat# 78442 |
| PageRuler Plus Prestained Protein ladder | Thermo Fisher Scientific | Cat# 26619 |
| Diphtheria Toxin | Millipore Sigma | Cat# D0564 |
| **Critical Commercial Assays** | | |
| Platinum II Host-Start PCR Master Mix (2x) | Thermo Fisher Scientific | Cat# 14000012 |
| PureLink RNA Mini Kit | Thermo Fisher Scientific | Cat# 12183025 |
| Maxima H-Minus First Strand cDNA Synthesis Kit with dsDNAse | Thermo Fisher Scientific | Cat# K1682 |
| TAQMAN Fast Advanced Master Mix | Thermo Fisher Scientific | Cat# 4444557 |
| EasySep Release Mouse Pan-B Cell Isolation Kit | STEMCELL Technologies | Cat# 19844 |
| EasySep Release Mouse CD138 Positive Selection Kit | STEMCELL Technologies | Cat# 100-0601 |
| 1X TMB | Thermo Fisher Scientific | Cat# 00-4201-56 |
| Novex ECL Chemiluminescent Substrate Reagent Kit | Thermo Fisher Scientific | Cat# WP20005 |
| AEC Substrate Set | BD Biosciences | Cat# 551951; RRID: AB_2868954 |
| eBioscience Fixable Viability (Live-Dead) Dye eFluor 780 | Thermo Fisher Scientific | Cat# 65-0865-14 |
| Cell Extraction Buffer | Thermo Fisher Scientific | Cat# FNN0011 |
| Bolt Bis-Tris Mini Protein Gels, 4-12%, 1.0 mm | Thermo Fisher Scientific | Cat# NW04122BOX |
| Invitrogen PVDF/Filter Paper Sandwiches, 0.45 μm | Thermo Fisher Scientific | Cat# LC2005 |
| **Experimental Models: Organisms/Strains** | | |
| Mouse: Jchain-DTR mice (J-DTR) | This paper | Not applicable |
| **Oligonucleotides** | | |
| *DTR* Genotyping WT Forward Primer:  GTCAAGTATTCCTTGCTGTGCAGATGATTAGG | This paper | Not applicable |
| *DTR* Genotyping Mutant Forward Primer:  GGTTACCATGGAGAGAGGTGT | This paper | Not applicable |
| *DTR* Genotyping Shared Reverse Primer:  ACTTCTGGGTGCAAATGGAGA | This paper | Not applicable |
| *FLP* Genotyping *FLP* Forward Primer:  ACAGAGACAAAGACAAGCGTTAGTAGG | This paper | Not applicable |
| *FLP* Genotyping *FLP* Reverse Primer:  ATTTCCCACAACATTAGTCAACTCCGTTAGG | This paper | Not applicable |
| *FLP* Genotyping Control DNA Forward Primer:  CTGCAACTCCAGTCTTTCTAGAAGATG | This paper | Not applicable |
| *FLP* Genotyping Control DNA Reverse Primer:  CCAGCTACAGCCTCGATTTGTGGTG | This paper | Not applicable |
| TaqMan Mouse *Actb* (Mm01205647_g1) | Thermo Fisher Scientific | Cat# 4331182 |
| TaqMan Mouse *Prdm1* (Mm00476128_m1) | Thermo Fisher Scientific | Cat# 4331182 |
| TaqMan Mouse *Jchain* (Mm00461780_m1) | Thermo Fisher Scientific | Cat# 4331182 |
| TaqMan Human *HBEGF (DTR)* (Hs00961129_m1) | Thermo Fisher Scientific | Cat# 4331182 |
| **Software and Algorithms** | | |
| FlowJo (v10) | BD Biosciences | RRID: SCR_008520 |
| GraphPad Prism 8 | GraphPad Software | RRID: SCR_002798 |
| Adobe Illustrator | Adobe | RRID: SCR_010279 |
| Adobe Photoshop | Adobe | RRID: SCR_014199 |
| BioRender | BioRender | RRID: SCR_018361 |
