## Supplementary material for "Jchain-Diphtheria Toxin Receptor Mice Allow for Diphtheria Toxin-Mediated Depletion of Antibody-Secreting Cells and Analysis of Differentiation Kinetics": Figures S1-S8

### Figure S1

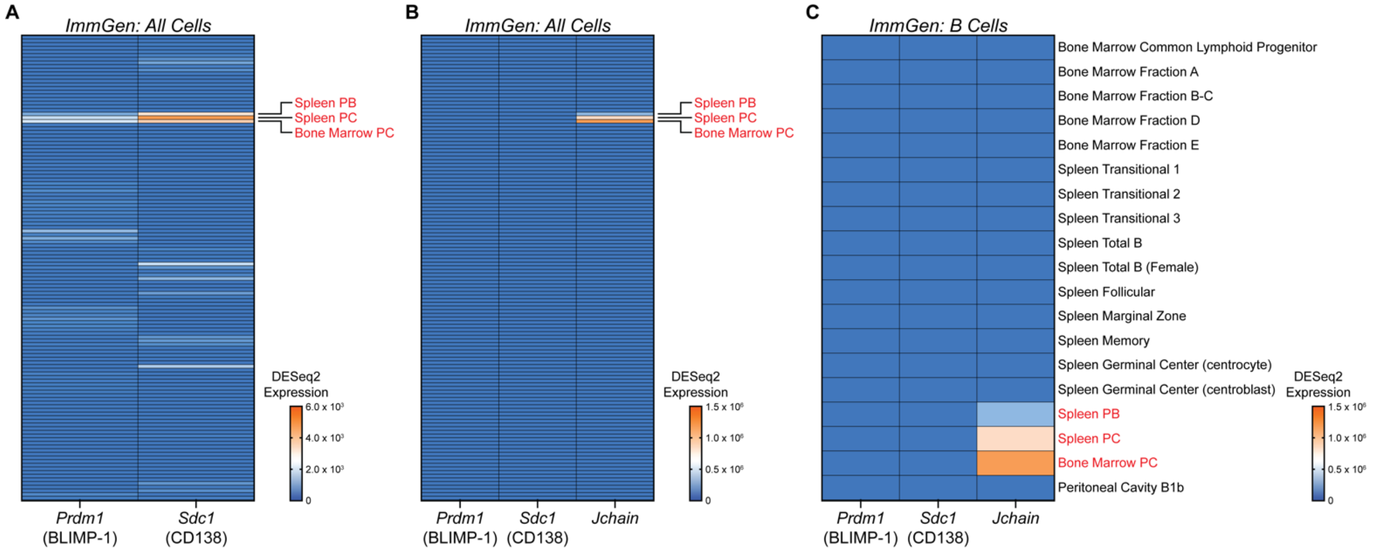


**Figure S1: *Jchain* is highly expressed in ASCs. (A)** ImmGen data showing *Prdm1* and *Sdc1* gene expression for all cell types. **(B)** ImmGen data showing *Prdm1*, *Sdc1* and *Jchain* gene expression for all cell types. **(C)** ImmGen data showing *Prdm1*, *Sdc1* and *Jchain* gene expression for all B cell subsets. **(A-C)** Data are derived from the ImmGen RNA-sequencing database and shown as DESeq2 processed expression levels.

### Figure S2

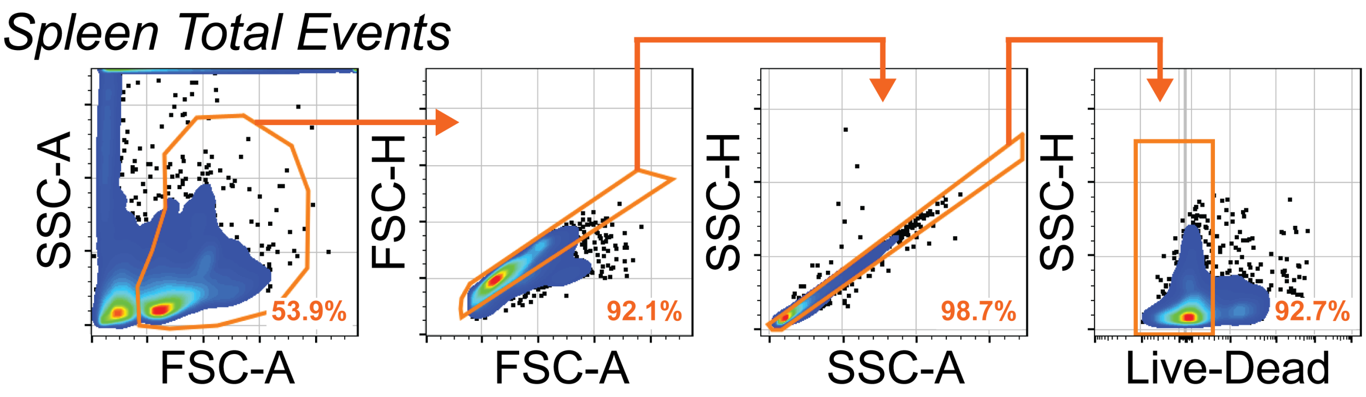


**Figure S2: Singlet and live cell gating strategy.** Representative gating of singlets and live cells using spleen as an example. The same strategy was applied to all flow cytometric analysis. Numbers in plots indicate percentages of gated populations within the immediate parent population.

### Figure S3

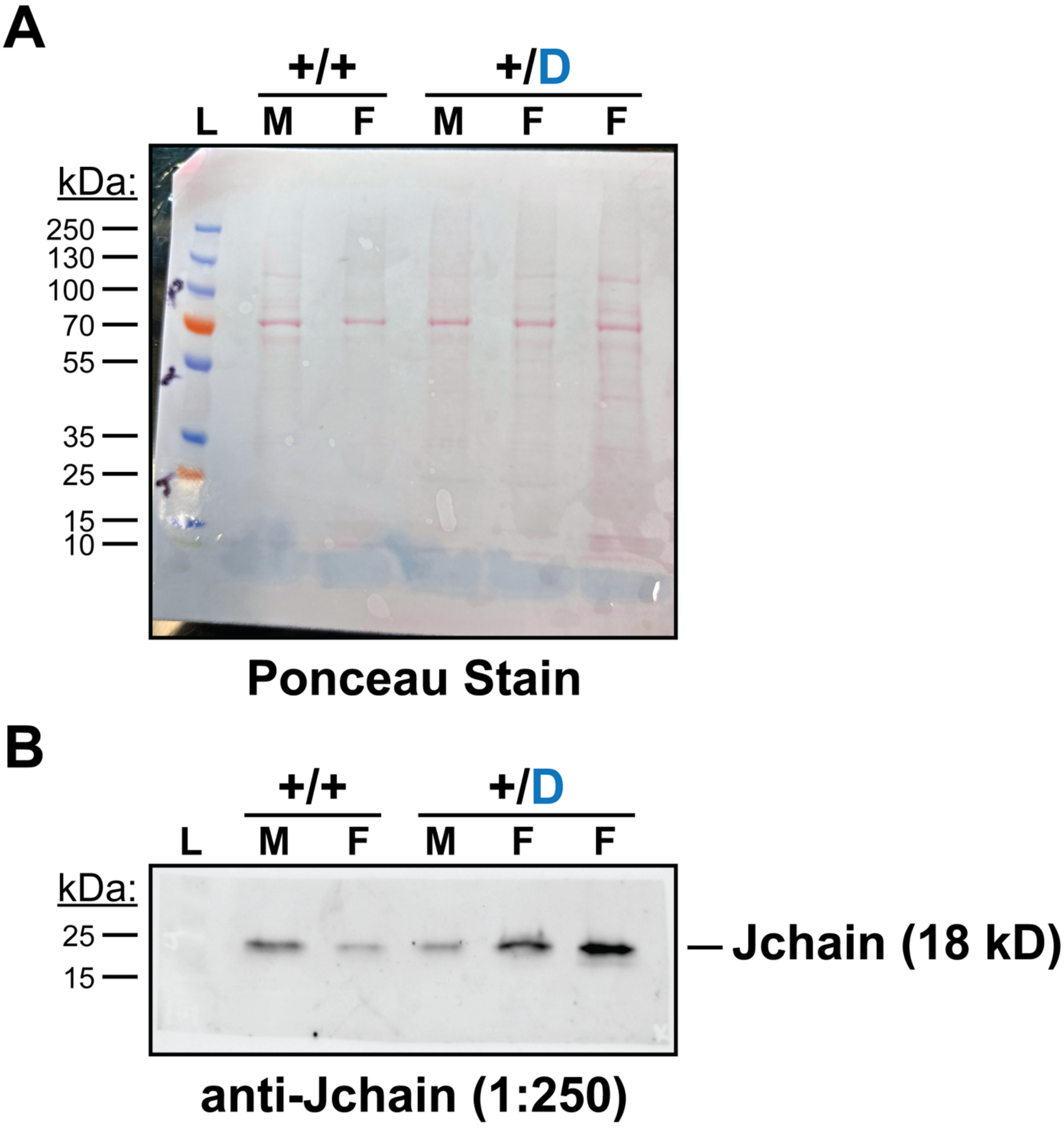


**Figure S3: Jchain protein is expressed in J-DTR spleen ASC-enriched samples. (A)** Ponceau stained membrane showing protein loading for spleen ASC-enriched samples from female (F) and male (M) WT (+/+) and J-DTR (+/D) mice. Molecular weights for the PageRuler Plus Prestained Protein Ladder (L) are provided for reference. Auto Tone in Photoshop was applied to the Ponceau stained membrane for visualization. **(B)** Western blot detection of Jchain protein for spleen ASC-enriched samples. The bottom third of the membrane from **(A)** was cut away and separately probed with an anti-Jchain primary antibody diluted 1:250. Subsequently, the membrane was probed with a goat anti-rabbit IgG-HRP secondary antibody diluted at 1:10,000. The membrane was imaged on a Bio-Rad ChemiDoc using chemiluminescence detection and is presented as is.

### Figure S4

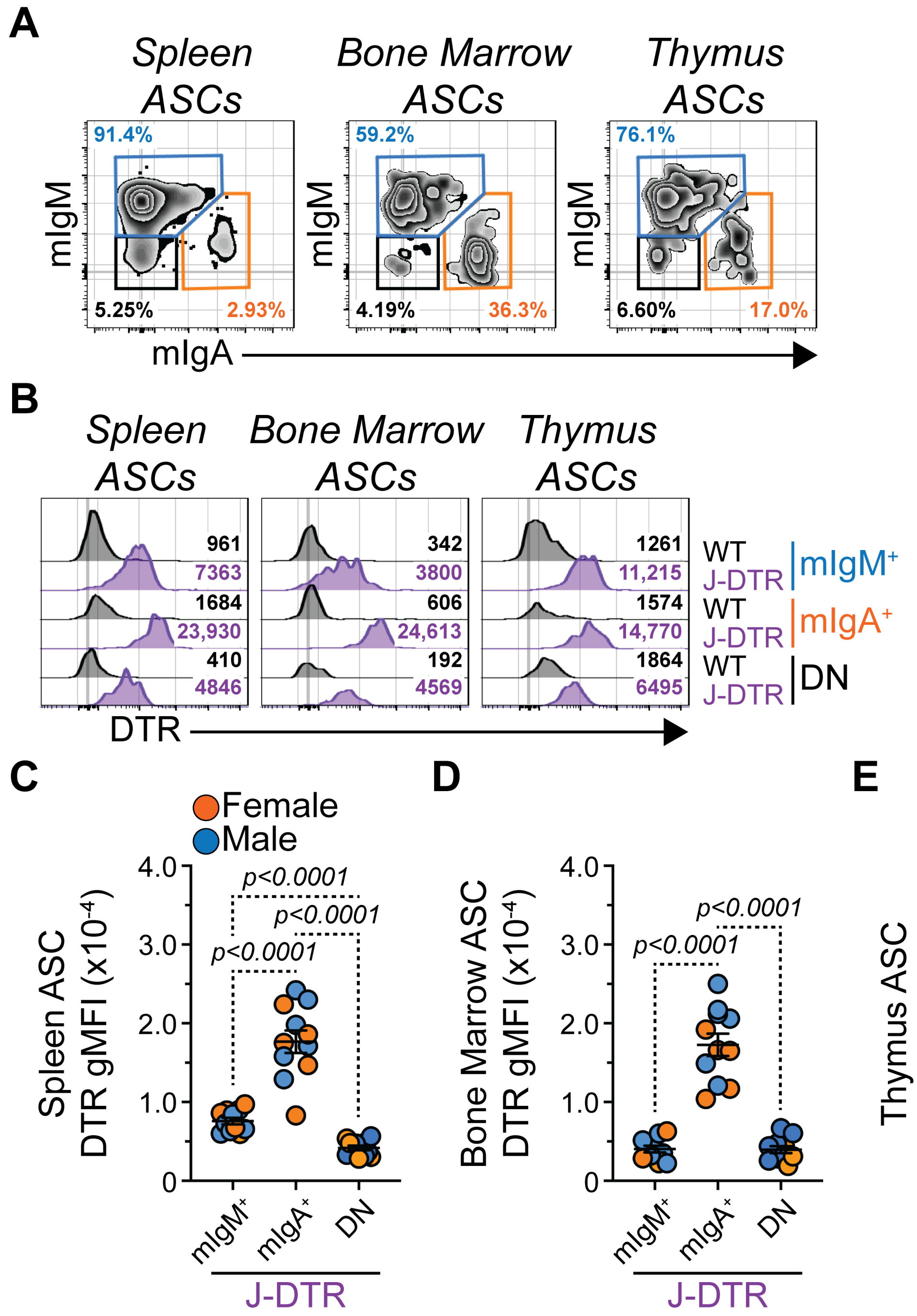


**Figure S4: Validation of DTR surface protein expression by mIgM$^+$, mIgA$^+$ and DN ASCs from J-DTR mice.** **(A)** Representative flow cytometry zebra plots showing gating of mIgM$^+$, mIgA$^+$ and DN ASCs from spleen, bone marrow and thymus. Numbers in plots indicate percentages of gated populations within the immediate parent population. **(B)** Representative flow cytometry histogram overlays showing surface expression of DTR by mIgM$^+$, mIgA$^+$ and DN ASCs from both WT and J-DTR mice. Data from spleen, bone marrow and thymus are shown. Numbers in plots indicate DTR gMFIs. **(C-E)** DTR gMFIs for WT and J-DTR mIgM$^+$, mIgA$^+$ and DN ASCs from **(C)** spleen, **(D)** bone marrow and **(E)** thymus. Symbols represent individual 3-5 months old female (orange) and male (blue) J-DTR mice. Horizontal lines represent mean ± SEM. J-DTR: female n = 5, male n = 6. Statistics: One-way ANOVA with Tukey's multiple comparisons test between ASC populations.

### Figure S5

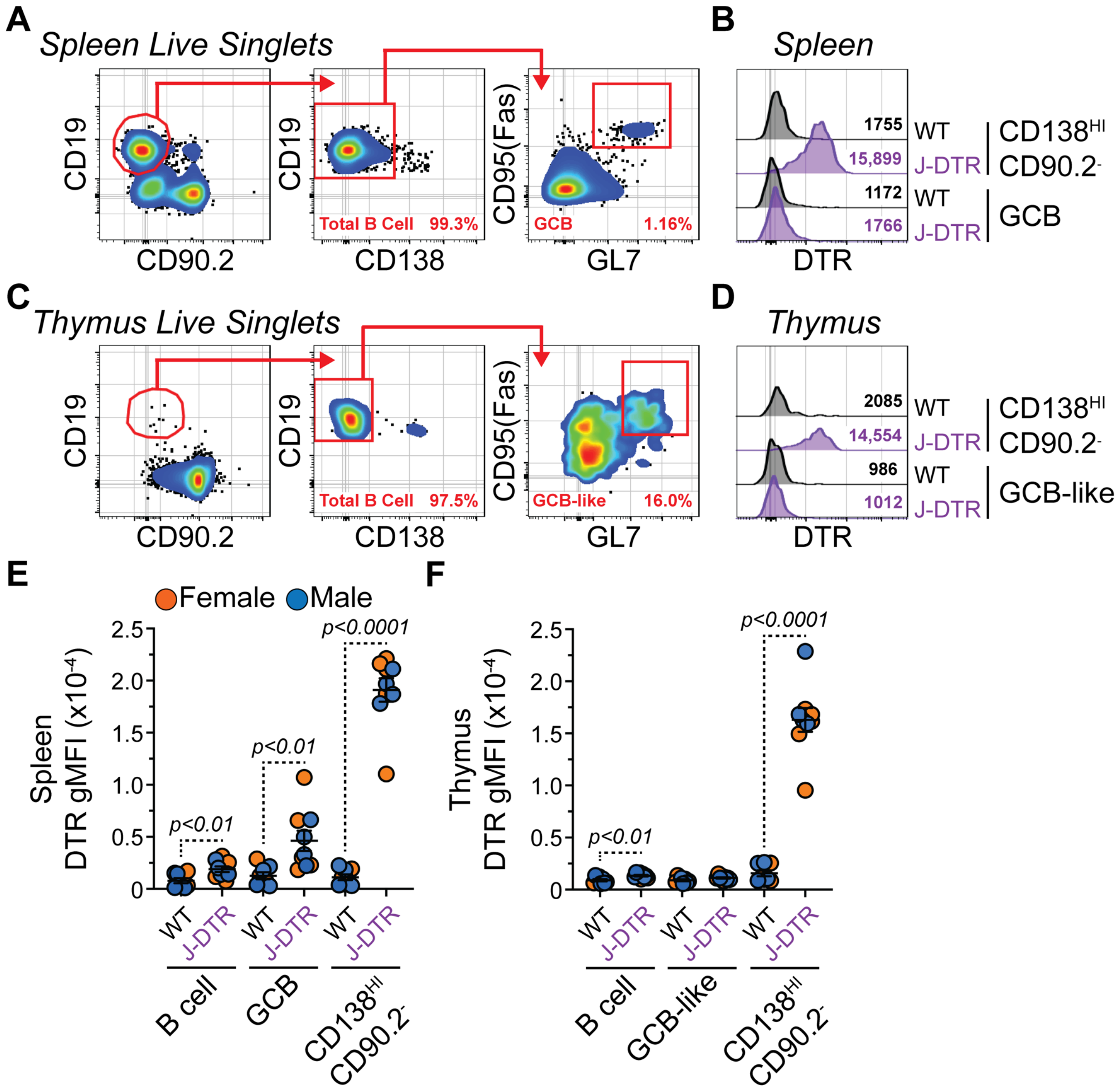


**Figure S5: DTR surface protein expression by B cells from J-DTR mice. (A)** Representative flow cytometry pseudocolor plots showing gating of spleen B cells and GCBs. Numbers in plots indicate percentages of gated populations within the immediate parent population. **(B)** Representative flow cytometry histogram overlays showing surface expression of DTR by spleen CD138$^{HI}$ CD90.2$^{-}$ cells and GCBs from both WT and J-DTR mice. Numbers in plots indicate DTR gMFIs. **(C)** Representative flow cytometry pseudocolor plots showing gating of thymus B cells and GCB-like cells. Numbers in plots indicate percentages of gated populations within the immediate parent population. **(D)** Representative flow cytometry histogram overlays showing surface expression of DTR by thymus CD138$^{HI}$ CD90.2$^{-}$ cells and GCB-like cells from both WT and J-DTR mice. Numbers in plots indicate DTR gMFIs. **(E-F)** DTR gMFIs for WT and J-DTR B cells, GCB (or GCB-like) and CD138$^{HI}$ CD90.2$^{-}$ cells from **(E)** spleen and **(F)** thymus. Symbols represent individual 3-7 months old female (orange) and male (blue) mice. Horizontal lines represent mean ± SEM. WT: female n = 4, male n = 4; J-DTR: female n = 5, male n = 4. Statistics: Unpaired Student's t-test comparing WT and J-DTR samples.

### Figure S6

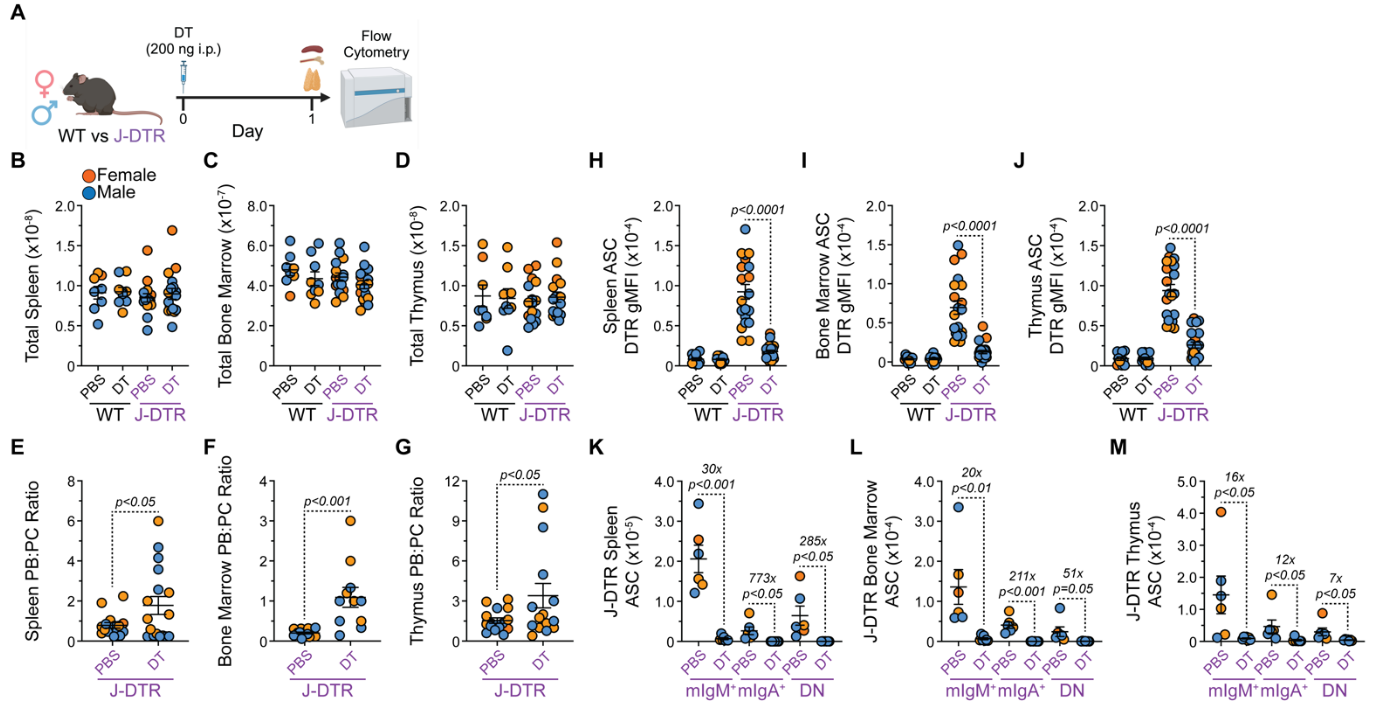


**Figure S6: Single dose administration of DT leads to the acute depletion of mIgM$^+$, mIgA$^+$ and DN ASCs in J-DTR mice. (A)** Schematic showing DT treatment of WT and J-DTR mice. 3-4 months old animals were given a single i.p. dose of 200 ng DT in 100 µL 1x PBS. Control mice received 100 µL of 1x PBS. Mice were euthanized the next day and spleen, bone marrow and thymus were assessed for ASC populations via flow cytometry. Schematic made with BioRender. **(B-D)** Total cell numbers for **(B)** spleen, **(C)** bone marrow and **(D)** thymus of WT and J-DTR mice treated with PBS or DT. **(E-G)** PB:PC ratios from **(E)** spleen, **(F)** bone marrow and **(G)** thymus of J-DTR mice treated with PBS or DT. Ratios calculated using absolute cell numbers. Only animals with non-zero populations for both PBs and PCs were assessed. **(H-J)** DTR gMFIs for ASCs from **(H)** spleen, **(I)** bone marrow and **(J)** thymus of WT and J-DTR mice treated with PBS or DT. **(K-M)** Numbers of mIgM$^+$, mIgA$^+$ and DN ASCs from **(K)** spleen, **(L)** bone marrow and **(M)** thymus of J-DTR mice treated with PBS or DT. **(B-M)** Symbols represent individual female (orange) and male (blue) mice. Horizontal lines represent mean ± SEM. Statistics: Unpaired Student's t-test with comparisons made between PBS and DT treatments within a genotype. **(B-D)** WT PBS: female n = 3, male n = 5; WT DT: female n = 4, male n = 5; J-DTR PBS: female n = 8, male n = 8; J-DTR DT: female n = 8, male n = 9. **(E-G)** J-DTR PBS: female n = 8, male n = 8; J-DTR DT: female n = 8, male n = 9. **(H-J)** WT PBS: female n = 3, male n = 8; WT DT: female n = 5, male n = 8; J-DTR PBS: female n = 10, male n = 10; J-DTR DT: female n = 10, male n = 11. **(K-M)** J-DTR PBS: female n = 3, male n = 3; J-DTR DT: female n = 3, male n = 4.

### Figure S7

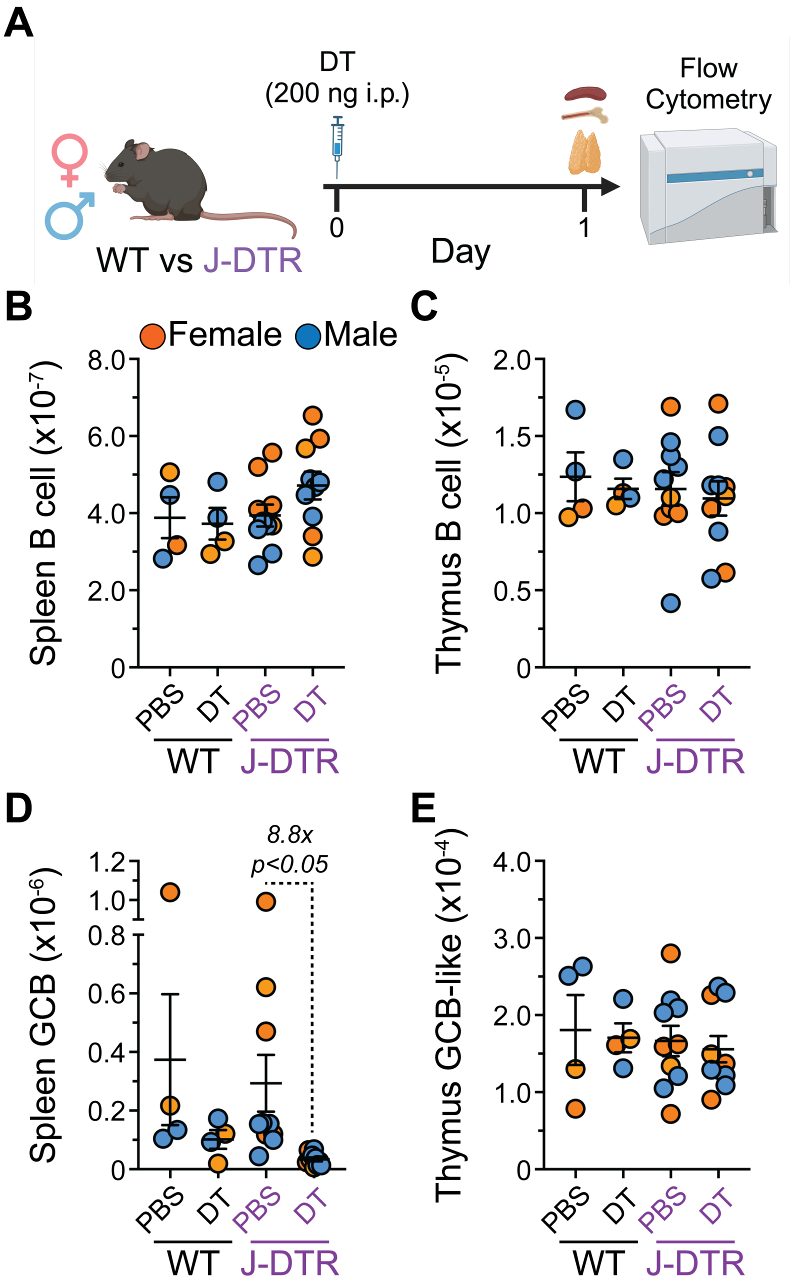


**Figure S7: Single dose DT administration leads to a modest spleen GCB reduction in J-DTR mice. (A)** Schematic showing DT treatment of WT and J-DTR mice. 3-4 months old animals were given a single i.p. dose of 200 ng DT in 100 µL 1x PBS. Control mice received 100 µL of 1x PBS. Mice were euthanized the next day and spleen and thymus B cell populations were assessed via flow cytometry. Schematic made with BioRender. **(B-C)** B cell numbers for **(B)** spleen and **(C)** thymus of WT and J-DTR mice treated with PBS or DT. **(D-E)** GCB (or GCB-like) numbers for **(D)** spleen and **(E)** thymus of WT and J-DTR mice treated with PBS or DT. **(B-E)** Symbols represent individual female (orange) and male (blue) mice. Horizontal lines represent mean ± SEM. WT PBS and DT: female n = 2, male n = 2; J-DTR PBS and DT: female n = 5, male n = 5. Statistics: Unpaired Student's t-test with comparisons made between PBS and DT treatments within a genotype.

### Figure S8

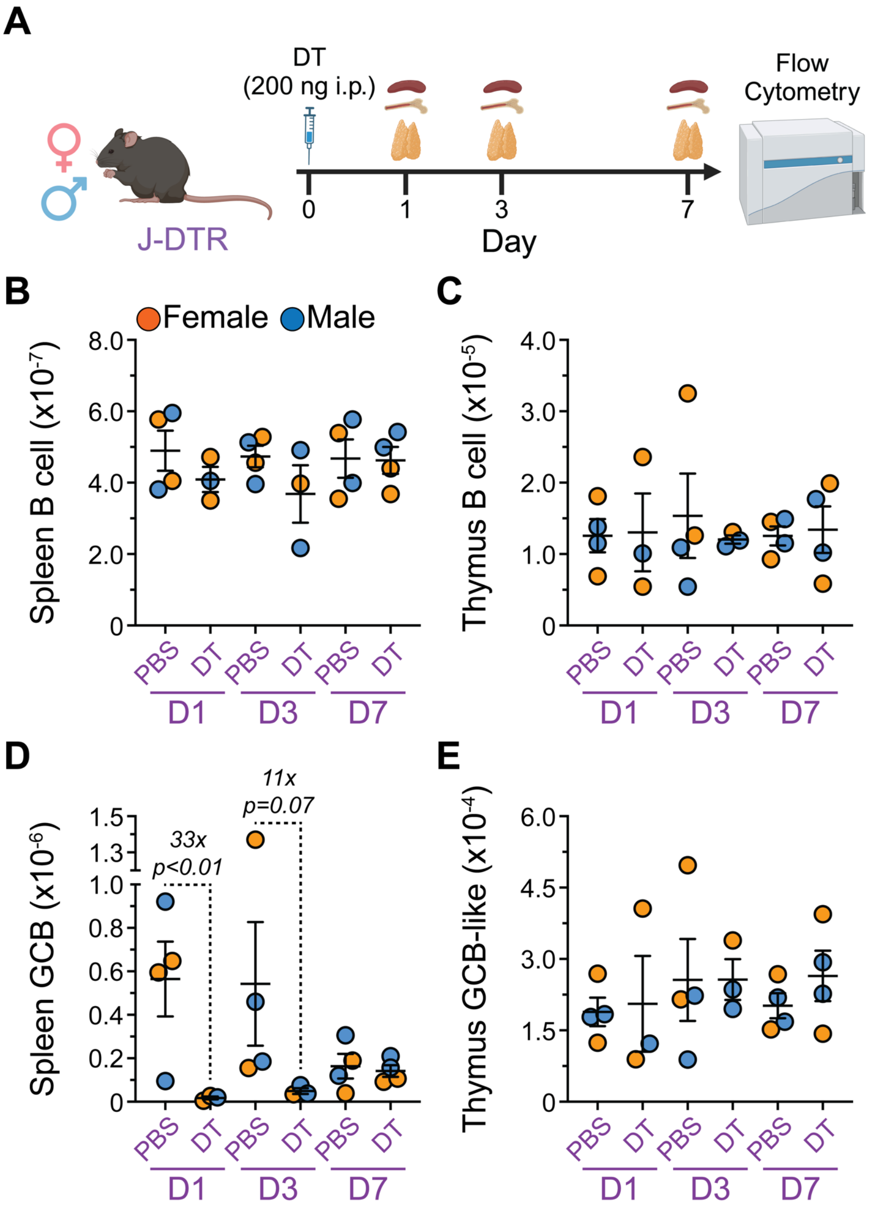


**Figure S8: Single dose DT administration results in transiently reduced spleen GCBs in J-DTR mice. (A)** Schematic showing DT treatment of J-DTR mice. 3-4 months old mice were given a single i.p. dose of 200 ng DT in 100 μL 1x PBS. Control mice received 100 μL of 1x PBS. Mice were euthanized at days 1, 3 and 7 post-injection. Spleen, bone marrow and thymus were assessed for B cell populations via flow cytometry. Schematic made with BioRender. **(B-C)** B cell numbers for **(B)** spleen and **(C)** thymus of J-DTR mice treated with PBS or DT. **(D-E)** GCB (or GCB-like) numbers for **(D)** spleen and **(E)** thymus of J-DTR mice treated with PBS or DT. **(B-E)** Symbols represent individual female (orange) and male (blue) mice. Horizontal lines represent mean ± SEM. J-DTR day 1 PBS: female n = 2, male n = 2; J-DTR day 1 DT: female n = 2, male n = 1; J-DTR day 3 PBS: female n = 2, male n = 2; J-DTR day 3 DT: female n = 1, male n = 2; J-DTR day 7 PBS: female n = 2, male n = 2; J-DTR day 7 DT: female n = 2, male n = 2. Statistics: Kruskal-Wallis test (nonparametric) with Dunn's multiple comparisons test. Comparisons made between PBS and DT treatments for a given day.
